# The onset of whole-body regeneration in *Botryllus schlosseri*: morphological and molecular characterization

**DOI:** 10.1101/2022.01.04.472827

**Authors:** Lorenzo Ricci, Bastien Salmon, Caroline Olivier, Rita Andreoni-Pham, Ankita Chaurasia, Alexandre Alie, Stefano Tiozzo

**Affiliations:** Laboratoire de Biologie du Développement de Villefranche-sur-Mer (LBDV), CNRS, Sorbonne University, 06230 Paris, France; Department of Organismic and Evolutionary Biology, Harvard University, Cambridge, MA 02138, USA; Institute for Research on Cancer and Aging in Nice (IRCAN), CNRS, INSERM, Université Côte d’Azur, 06107 Nice, France

## Abstract

Colonial tunicates are the only chordates that regularly regenerate a fully functional whole body as part of their asexual life cycle, starting from specific epithelia and/or mesenchymal cells. In addition, in some species, whole-body regeneration (WBR) can also be triggered by extensive injuries, which deplete most of their tissues and organs and leave behind only small fragments of their body.

In this manuscript, we characterized the onset of WBR in *Botryllus schlosseri,* one colonial tunicate long used as a laboratory model. We first analyzed the transcriptomic response to a WBR-triggering injury. Then, through morphological characterization, *in vivo* observations via time-lapse, vital dyes, and cell transplant assays, we started to reconstruct the dynamics of the cells triggering regeneration, highlighting an interplay between mesenchymal and epithelial cells. The dynamics described here suggest that WBR in *B. schlosseri* is initiated by extravascular tissue fragments derived from the injured individuals rather than particular populations of blood-borne cells, as has been described in closely related species. The morphological and molecular datasets here reported provide the background for future mechanistic studies of the WBR ontogenesis in *B. schlosseri* and allow to compare it with other regenerative processes occurring in other tunicate species and possibly independently evolved.

## 1 Introduction

Within the lifespan of a metazoan, sub-lethal damages or loss of body parts can occur frequently as a consequence of predation, competition, pathogens infections, or simply by accident. Animals cope with such traumatic events by developing a wide range of strategies, such as the synthesis of protective structure, scar formation, or various degrees of regeneration (Vorontsova and Liosner, 1961; Sinigaglia et al. *in press*). The most extreme examples of regeneration occur when the entire functional body is restored from only minute fragments of the original organism, a *bona fide* ontogenesis generally referred to as whole-body regeneration (WBR). WBR has been described in many animal species belonging to different non-vertebrate taxa (Blanchoud and Galliot, 2022), and it is often correlated with the capacity of such organisms to reproduce asexually, i.e. a cyclical form of body regeneration that suggests possible co-options of cellular and molecular mechanisms between the injury-triggered and the physiological WBRs (Martinez et al., 2005; Sánchez Alvarado and Yamanaka, 2014; Sinigaglia et al. *in press*).

Colonial species of tunicates, the sister group of vertebrates, acquired the capacity to undergo WBR as part of their asexual life-cycle and also as a response to extreme injury, they are therefore promising models to compare these two forms of non-embryonic development, study their mechanisms and infer their evolution (Alié et al., 2020). During tunicate asexual reproduction, adult individuals called zooids regenerate cyclically the entire body through a non-embryonic developmental process generally called propagative budding (Nakauchi, 1982), ultimately leading to the formation of colonies of genetically identical individuals. The way the propagative budding processes unfold differs from one species to another, starting from different and often non-homologous cells and tissues, but often converging into a common stage of two concentric hollow vesicles, each of them formed of a monolayer epithelium, reviewed in (Alié et al., 2020). From this phylotypic asexual stage of double-vesicle, the process of organogenesis begins, eventually leading to a *bauplan* that is shared by the whole subphylum (Alié et al., 2020). In many tunicates, the capacity of WBR is not only a characteristic of their life-cycle, but it can also be triggered in response to extensive injury (Tiozzo et al., 2008a), in which case the WBR process is referred to as survival budding (Nakauchi, 1982).

Both propagative and survival buddings have been studied mainly in the subfamily of *Botryllinae* (Brunetti, 2009)(**Supplementary Figure 1**), a widespread group of colonial tunicates composed of small zooids (<0.5cm) embedded in a common soft extracellular matrix, the tunic, and connected by an extracorporeal network of the epidermal derived vessels (Manni et al., 2007; Tiozzo et al., 2008c). Throughout the vasculature, different types of mesenchymal cells, the hemocytes, circulate through the colony propelled by zooids’ hearts and by the peristaltic movement of ampullae, the blind tips of the circulatory vessels (**Supplementary Movie 1**). Botryllids include several species of the genera *Botryllus* and *Botrylloides,* which undergo WBR via two modes of budding: peribranchial budding, a form of propagative budding that arises from a multipotent epithelium(Manni et al., 2014; Ricci et al., 2016b; Alié et al., 2020), and vascular budding (VB) that, depending on the species, can be propagative or triggered by injury (Oka and Watanabe, 1957, 1959; Milkman, 1967; Satoh, 1994) (**Supplementary Figure 1**). For instance, in *Botryllus primigenus* VB occurs routinely in a propagative fashion, while in other botryllids, such as *Botrylloides violaceus* (Brown et al., 2009) and *Botrylloides leachi* (Rinkevich et al., 2007), VB occurs upon the exogenous removal of the existing zooids. It has then been suggested that the source of cells forming both propagative and survival vascular buds is a population of hemocytes that aggregate in the vascular network. Recently, Kassmer and collaborators (Kassmer et al., 2020) identified a population of Integrin-alpha-6-positive (Ia6+) hemocytes as candidate stem cells responsible for induced VB in the species *Botrylloides diegensis*. Ia6+ hemocytes, which constantly divide in healthy colonies, also express genes associated with pluripotency. The latter findings strongly suggest that the presence of permanent population/s of circulating stem cells may be at the bases of WBR via vascular budding in botryllid tunicates.

The species *Botryllus schlosseri* has been widely used in the last several decades as a laboratory model for developmental biology, immunology, and regenerative biology (Manni et al., 2007, 2019; Kürn et al., 2011; Voskoboynik and Weissman, 2014; Gasparini et al., 2015; Munday et al., 2015; Kassmer et al., 2016). In *B. schlosseri*, VB occurs purely in response to injury, and it can be triggered in laboratory conditions by depleting the colony of the adult zooids and their peribranchial buds via microsurgery (Milkman, 1967; Sabbadin et al., 1975; **Supplementary Figure 2A-B**). While asexual propagation through peribranchial budding has been increasingly characterized these past years (Tiozzo et al., 2005; Manni et al., 2014; Di Maio et al., 2015; Ricci et al., 2016a, 2016b; Pruenster et al., 2018; Prünster et al., 2019), only a few studies have addressed VB in this species (Milkman, 1967; Sabbadin et al., 1975; Voskoboynik et al., 2007; Ricci et al., 2016a; Nourizadeh et al., 2021). The cell populations and the tissues involved in the onset of *B. schlosseri* VB are still not well defined, and the morphogenetic events that lead to the regeneration of a functional adult zooid are poorly described. In this manuscript, we follow the dynamic of WBR upon injury in the laboratory model *Botryllus schlosseri*. We focus on the early stages of the process and characterize the transcriptome profile of the initial response of the whole colony to extensive injury; we describe the cytological and histological structures at the onset of the presumptive vascular bud and test the contribution of mesenchymal cells and vascular epithelia.

The correlated observations suggest that WBR is initiated by extravascular tissue fragments derived from the injured zooids or buds, rather than a particular population of hemocytes as occurring in other closely related species.

## 2 Material and Methods

### 2.1 Animal culturing and surgical procedure

Colonies of *Botryllus schlosseri* were raised on glass slides in a marine-culture system as described previously (Langenbacher et al., 2015). Colonies used for WBR induction experiments were transferred to an 18°C incubator in small containers (<1L) in a closed system with filtered seawater (FSW) and bubblers, with a day/night cycle of 10h/14h and no feeding. The water was completely replaced every two days. Colonies of *B. schlosseri* at stage D (Lauzon et al., 2002) were dissected with microsurgery tools and syringe needles (30G, Terumo, SG2-3013) under a stereomicroscope. After the removal of all zooids and peribranchial buds, animals were cleaned and allowed to regenerate in FSW. Water was replaced every two days and vascular bud detection was performed by daily observations under a stereomicroscope allowing a 120X magnification. For fluorescent *in situ* hybridization (FISH) experiments, dissected colonies were fragmented into small pieces before fixation to facilitate the penetration of solutions (Prünster et al., 2019).

### 2.2 Video acquisition and processing

Regenerating colonies were placed in a room at 18°C in a petri dish filled with 150ml of FSW. Photographs for time-lapse videos were taken every 5 min for up to 8 days post-injury using a Canon EOS 6D Mark II equipped with a 100mm macro objective. Videos were assembled using Avidemux 2.7.8 (http://www/avidemux.org).

### 2.3 Immunohistochemistry

Whole *B. schlosseri* systems and regenerating colonies were anesthetized in natural seawater and MS222 0,3% (Sigma-Aldrich, #E10505-25G) and processed as previously described (Ricci et al., 2016a). Nuclei were counterstained by incubation at room temperature with 1 μg/ml Hoechst 33342 in PBS for 2h, then mounted in glycerol after quick washes in PBS. Confocal were acquired with a Leica TCS SP5, SP8, or Stellaris microscope. Primary antibodies include: polyclonal, mouse anti-integrin-alpha 6 (DSHB, P2C62C4) diluted 1:10 in PBS; polyclonal, rabbit anti-phospho Histone H3 (Ser10), (Merk Millipore #06-570) diluted 1:1000 in PBS; monoclonal, mouse anti acetylated tubulin (Sigma-Aldrich #T6793), diluted 1:1000 in PBS; monoclonal, mouse anti tyrosinated tubulin, (Sigma-Aldrich #T90028), 1:1000; monoclonal, mouse anti-gamma tubulin (Sigma-Aldrich, # T6557) 1:500; polyclonal rabbit anti-PKCξ C-20 (Santa-Cruz Biotechnology inc., #sc-216) 1:1000; anti-phospho-tyrosine, 4G10® Platinum, (Merck Millipore, #05-1050X) 1:500.

### 2.4 Transmission electron microscopy (TEM)

Samples for ultra-thin sectioning were fixed with a 3% solution of glutaraldehyde in sodium cacodylate buffer (pH 7.3), post-fixed with osmium tetroxide (OsO4) 1% in cacodylate buffer, dehydrated using acetone, and embedded in epoxy resin. An UltracutE Reichert ultramicrotome was used for the ultra-thin sections (60–80 nm), which were contrasted with uranyl acetate and lead citrate and observed under a transmission electron microscope TEM JEM 1400 JEOL coupled with a MORADA SIS camera (Olympus).

### 2.5 Whole-mount in situ hybridization

Antisense mRNA probes were designed within the coding region of each gene (**Supplementary Figure 3**) FISH was carried out as previously described (Ricci et al., 2016a). DIG-probe detection was performed with bench-made FITC-Tyramide and TRITC-Tyramide by 3 hours incubation.

### 2.6 In vivo cells and tissue labeling and imaging

Colonies were grown in Willco-dishes (Willco-Dish®, 50x7x0.17mm). Once reached stage D (Lauzon et al., 2002) the colonies were injected with 1-2 µl per system of lipophilic dye FM® 4-64 Dye, (Life Technologies, #T-13320), diluted 1: 100 in PBS and with BSA Alexa Fluor® 488 conjugate, (Life Technologies, #A13100) at a concentration of 1mg/mL, according to published parameters (Braden et al., 2014). Following injection, colonies were left to recover 3hrs in FSW, then dissected to induce WBR, and left to regenerate in FSW. After vascular bud detection, the colony was observed with a confocal Leica TCS SP5 microscope vesicle.

### 2.7 Fusion-chimera assay and genotyping via microsatellite

To trigger fusion, two isogenic and histocompatible colonies were selected and sub-cloned side-by-side in the same glass slide. The colonies were allowed to grow until they fuse. Following fusion, hemocytes cells from both genotypes were immediately mixed in the plasma and circulated freely in the whole vascular system of the chimera. Around 48h after fusion, the couple of colonies were separated, and, as soon as they reached stage D, WBR was induced as previously described (**Supplementary Figure 2C**). For micro-satellite sequencing, after fusion of allogeneic colonies, clear landmarks were established to delineate the vascular system of each colony by scratching the glass slide with a diamond pen and taking photographs of the colony prior and daily after fusion. Dissected colonies were left to regenerate until they produced a vascular bud that underwent organogenesis. Large vascular buds were dissected with microsurgery tools and syringe needles (30G, Terumo, SG2-3013) under a stereomicroscope. Stomach epithelium was isolated with thin forceps and repetitively washed in clean FSW to avoid blood cell contamination. Then, genomic DNA was extracted from the stomach tissue, using the NucleoSpin® Tissue XS kit for genomic DNA (Mascherey-Nagel, #740901.50) and eluted in 10 µl of elution buffer. Following elution, samples were stored at -20°C. Tissues from both fused colonies were collected separately before fusion and their genomic DNA was collected with the same procedure as used for vascular buds. Couples of forward and reverse primers complementary to a *Botryllus* non-coding genomic locus designed to amplify microsatellites sequences BS1 and PB49 were used (Stoner and Weissman, 1996; Ben-Shlomo et al., 2008). For each microsatellite locus, a 5’ tag made of a universal oligo was added to the forward primer. The sequence of this universal primer was used to design another forward primer, with a 6-FAM™ fluorescent tag at its 5’ end (Life Technologies). Three primers PCR amplification were performed using the Qiagen Multiplex PCR kit (206143) in a final volume of 20 µl, at the concentration of 0.01 µM of forward primer and 0.2 µM of each reverse and 6-FAM forward primer. 1µl of gDNA was added to the reaction as a template. The cycling program was as follows: denaturation, 95°C, 15 min; amplification,( 94°C, 30s; 60°C, 90s; 72°C, 60s)x40; 60°C, 30 min for the BS1 locus. For the PB49 locus, the program was modified as follows: denaturation, 95°C, 15min; amplification, (94°C, 30s; 65°C, 1min; 72°C, 1min)x3 then (94°C, 30s; 63°C, 1min; 72°C, 1min;)x17 and (94°C, 30s; 57°C, 1min; 72°C, 1min)x20; 60°C, 30 min. The success of the PCR was validated by electrophoresis on a 1.7%, agarose gel before genotyping. Genotyping was performed by the Plateforme Génome Transcriptome de Bordeaux, Site de Pierroton – INRA. Three primers PCR products were diluted in formamide to avoid excessive fluorescence, with a dilution factor of 50 or 100, according to the sample. They were subsequently analyzed with an ABI3730 analyzer, in parallel with LIZ-600 and LIZ-1200 size standards. Fragments sizes were then analyzed with the Peak Scanner™ Software v2.0 (Applied Biosystems).

### 2.8 RNA extraction and Transcriptome sequencing and differential expression analyses (RNAseq-DEA)

An isogenic strain of *Botryllus schlosseri* was tested for its ability to regenerate and produce vascular buds in an average time of 2-3 days. Twelve subclones of comparable size from this strain were separated with a razor blade and allowed to grow separately on individual glass slides. When colonies reached stage D (Lauzon et al., 2002), they were dissected with microsurgery tools and syringe needles (30G, Terumo, SG2-3013) under a stereomicroscope. After removal of all zooids and buds, animals were cleaned and either conditioned for further RNA extraction or allowed to regenerate in Filtered seawater (FSW) in small containers (<1L) placed in an incubator at 19°C. When allowed to regenerate, colonies were left 6, 18, or 24 hours post-injury (hpi) in FSW before preparing for RNA extraction. The regenerating colony was detached from the glass slide with a razor blade and then transferred to a tube and flash frozen before storage at -80°C and later RNA extraction. For each time point, three replicates were made, bringing the total number of samples to twelve (**Figure 1A**).

**Figure 1.**
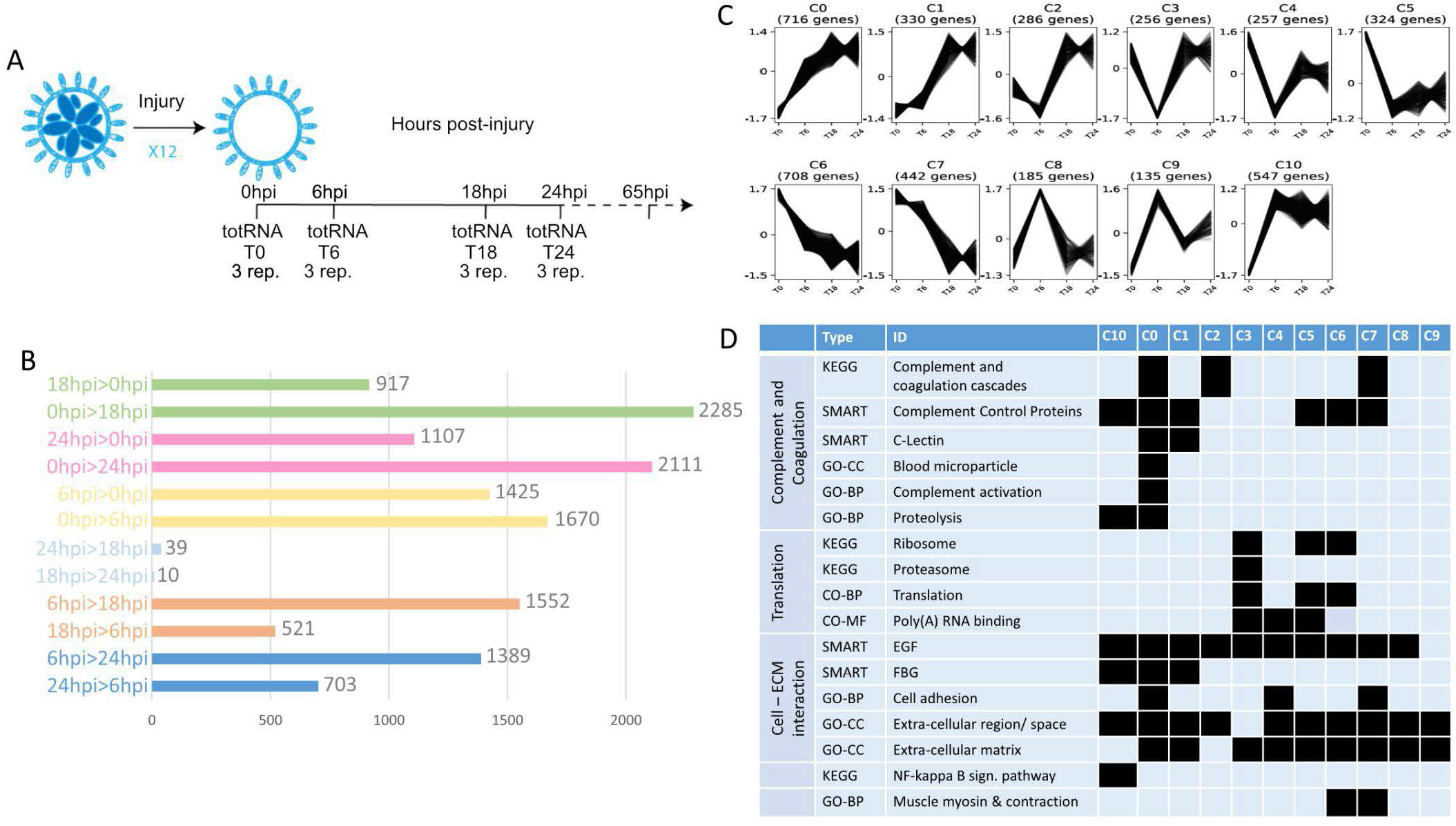
Transcriptomic profile of early steps of WBR in *B. schlosseri*. (A) Scheme showing the experiment design for transcriptome characterization of early WBR steps in *B. schlosseri*. In addition to the harvested colonies, additional clones were allowed to regenerate for 65hpi to certify the WBR ability of the clone. (B) Graph showing the number of contigs differentially expressed between each condition. (C) The eleven clusters of expression profile and the number of contigs in each cluster (D) Chosen examples of functional categories statistically enriched across the eleven clusters, extracted from Supplementary Table 1. GO-CC, Gene Ontology - Cell Compartment; GO-BP, Gene Ontology - Biological Process; GO-MF, Gene Ontology - Molecular Function.

Extraction of total RNA was performed the same day in a single round, for the twelve samples, using the NucleoSpin® RNA XS Mascherey-Nagel kit (#740902.50). First, 500μl of lysis buffer from the kit was added to the 1.5 ml tubes containing the samples. The latter was subsequently ground manually in the tube, using a plastic, RNAse free micropestle. All further steps were performed according to the user manual section for RNA extraction from animal tissue. For each sample, total RNA was eluted in 12µm of nuclease-free water and stored at -80°C until sequencing.

Library preparation and sequencing were performed at the USC Epigenomic Center (Los Angeles, USA) according to the Illumina HiSeq 2500 protocol. Approximately 70M PE reads were sequenced for each of the twelve samples.

Transcriptome assembly and differential expression analysis were performed as follows. Step 1: removing of contaminating ribosomal RNA using SortmeRNA v2.1 (Kopylova et al., 2012); Step 2: cleaning, clipping, and filtering reads using Trimmomatic v0.30 (Bolger et al., 2014); Step 3: transcriptome assembly from the remaining reads, using Trinity v2.11.0 (Grabherr et al., 2011) with default parameters; Step 4: Recover the best ORF per contig using TransDecoder v5.5.0, using a minimum protein length of 90 amino-acids (-m parameter); Step 5: Reduce spurious redundancy by collapsing similar transcripts using cd-hit-est v4.6 with default parameters (Fu et al., 2012); Step 6: Mapping the sequencing reads on the obtained transcriptome using Kallisto v0.43.1 using default parameters (Bray et al., 2016); Step 7: Identifying Differentially Expressed Genes using DeSeq2 (Love et al., 2014) through the iDEP v0.93 platform, following a between-sample normalization of expression values given by Kallisto to ensure a homogeneous distribution of expression data across samples (see **Supplementary Table 1**).

Genes being differentially expressed between at least two conditions (e-value < 0.05 and Fold-change >1) have been clustered by expression profile using Clust v1.12.0 (Abu-Jamous and Kelly, 2018)using raw ESTs (Kallisto output) as input data. Then contigs were named after their best tblastn hit against the human uniProt_proteome_UP000005640. Functional enrichment for each cluster was investigated using DAVID Bioinformatics Resources 6.8, using the whole transcriptome (from Step 5 above) as a reference dataset.

## 3 Results

### 3.1 Wound healing response and vascular remodeling precede injury activated WBR

In order to describe the transcriptomic response to extensive colony injury, colonies of *Botryllus schlosseri* were allowed to regenerate for 0, 6, 18 and 24 hours post-injury (hpi) respectively (see Material and Methods, **Figure 1A**). Approximately 147 million reads were cleaned and assembled into 157,306 contigs (N50=707 nuc.) from which 32,561 open reading frames were retained for downstream analyses (**Supplementary Data 1**). Gene expression level across the four time-points was measured by mapping reads to the 32,561 contigs, leading to the identification of 6,007 contigs having a differential expression (adjusted p-value < 0.05 and Log2 Fold-change > 1) between at least two conditions (**Supplementary Table 1**, **Figure 1B**). Most of the variation in gene expression arises between 0, 6 and 18/24hpi, while 18hpi and 24hpi have similar molecular profiles (**Figure 1B**). The 6,007 contigs were grouped into eleven clusters based on their expression profiles (**Figure 1C**). Clusters 0, 1 and 10 correspond to a general increase in expression upon surgery; clusters 5, 6, and 7 to a general decrease, while the other clusters show more complex profiles (**Figure 1C**). Taken together these results show a drastic transcriptomic response to injury in the first 18 hours of WBR in *Botryllus*.

Functional enrichment of the retrieved clusters (**Supplementary Table 1**, **Figure 1D**) reflects the active role played by the circulatory system in injury response and WBR initiation, in line with the important vascular remodeling observed after zooid ablation (**Supplementary Movie 2**). Indeed, seven clusters are enriched in genes of the complement and coagulation cascade (**Figure 1D**), a mammalian proteolytic cascade in blood plasma acting as a defense mechanism against pathogens. More specifically, clusters 0, 1, 2 and 10 comprise orthologues of the complement components C3/C5 - the core proteins of the complement cascade - as well as transcripts similar to MASP and Ficolins that activate the complement through the lectin pathway (**Supplementary Table 1**, **Supplementary Figure 4**, **Supplementary Data 2**). Clusters 5, 6 and 7 contain genes involved in coagulation (*e.g.* orthologue or Coagulation factor XIII B chain) and platelet activation (*e.g.* selectin-like genes). The enrichment of Fibrinogen (FBG) domain-containing genes (clusters 0, 1, 10) and of ECM components (all clusters) (**Figure 1D**) suggests a link between blood clotting and vascular remodeling by modulation of the physical interactions between vascular epithelium and extracellular matrix. Putative regulators of angiogenesis (the formation of new vessels from pre-existing ones) are numerous in clusters 0 and/or 1 (**Supplementary Table 1**), including transcripts similar to tenascins and angiopoietins, as well as orthologues of the Angiopoietin receptor (TIE1/2) and the transcription factors ETS1 and Sox7/17/18 (**Supplementary Data 2**). In mammals, ETS-1 controls endothelial cell migration and invasion (Iwasaka et al., 1996), while Sox17 promotes angiogenesis and endothelium regeneration (Liu et al., 2019). Finally, *Botryllus schlosseri* Gata-b, the orthologue of Gata1/2/3 that we previously found expressed in vascular buds (Ricci et al., 2016a), also belongs to cluster 0. In mammals, Gata-2 is central to maintaining endothelial cell identity (Kanki et al., 2011).

Functional enrichment analysis also revealed an expression increase of the NF-kappa B signaling pathway, involved in mammalian immunity and cell survival (Oeckinghaus et al., 2011), as well as a drop in the expression of translation-related genes, especially of ribosomal protein-coding genes (**Supplementary Table 1**). The biological significance of the latter is still unclear, but it may be linked to the translational response to stress (Advani and Ivanov, 2019).

### 3.2 WBR origins from extravascular tissues that migrate into the vasculature

To track the origin of WBR, we filmed with a high-resolution camera the entire colonies of *B. schlosseri* upon microsurgery (n=9 colonies) and allowed them to regenerate until the morphogenesis of new zooids. The analyses of the digitally magnified areas of budding showed that, in the tunic near the dissection area, relatively small (50-70µm) fragments of tissues start to move towards the vasculature, get surrounded by the latter, and eventually develop into a new zooid (**Supplementary Movie 3-5**). Such tissue fragments are not present in the tunic of undissected colonies suggesting that they may be debris of zooids or peribranchial buds, left behind after dissection. To better understand the dynamic of WBR, we followed *in vivo* the migrating tissues within the tunic until they got in contact with the vasculature. Then, we fixed and examined the details of the tissue interactions (**Figure 2**). From the observation of n=11 putative WBR onsets from four different colonies, we detected the presence inside the tunic of double monolayered vesicles approaching (**Figure 2A-E’’, Supplementary Movie 6**) and fusing to (**Figure 2F-Q’’, Supplementary Movie 7-8**) the vasculature. We also reported more complex epithelial structures already fused to the vasculature (n=1) (**Figure 2R-U’’’**, **Supplementary Movie 9**). The presence of such intravascular structures has never been observed in undissected colonies during their asexual growth (data not shown).

**Figure 2.**
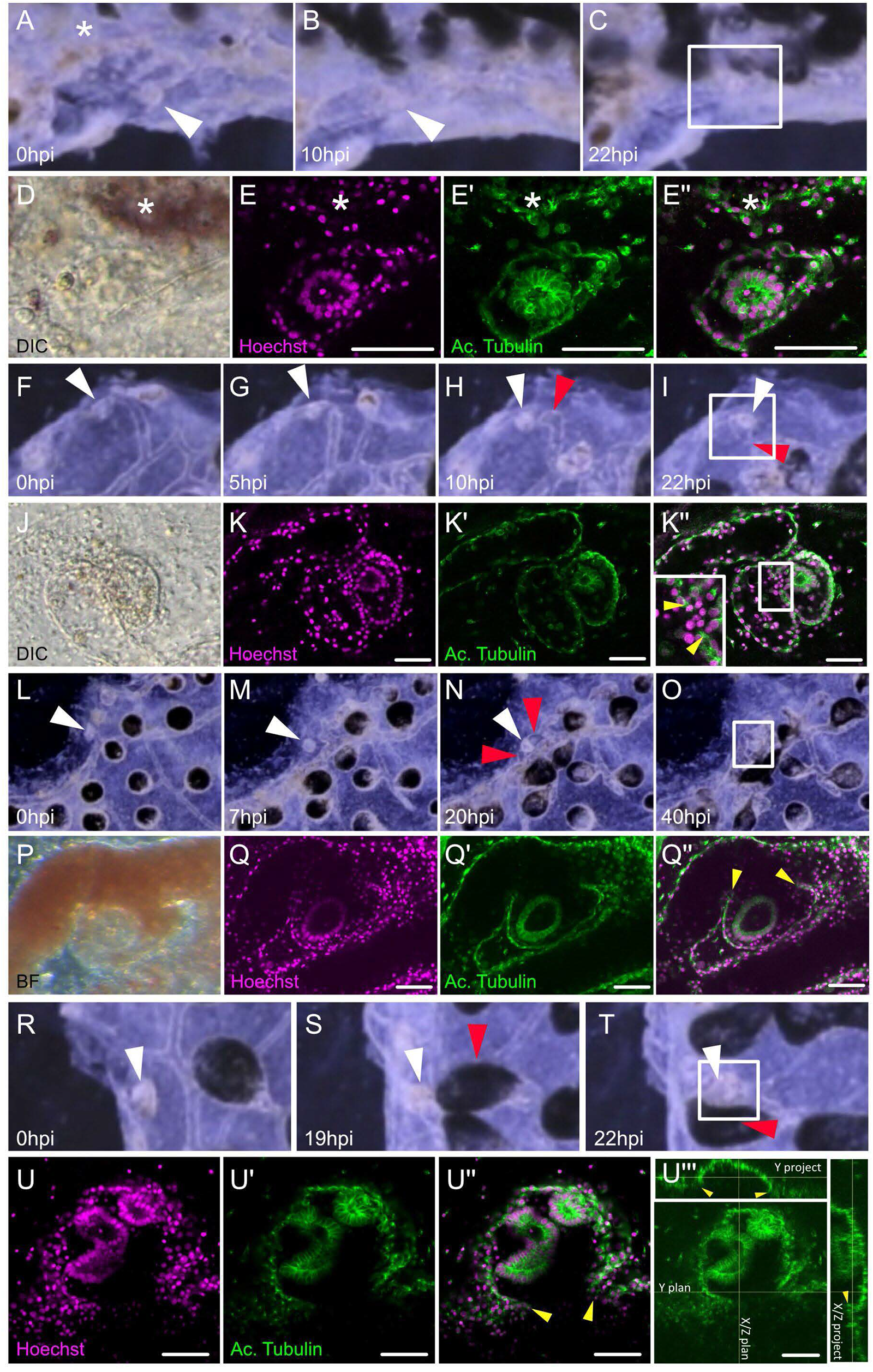
Dynamic of the migration of extravascular tissues into the vascular network. (A-C) Screenshots from Supplementary Movie 6 show tissue left-over getting in close contact to the vasculature from (A) 0 hours post-injury (hpi) to (B) 10hpi and (C) 22hpi. (D-E’’) Microscopic view of the areas squared in (C). (D) Transmitted light with DIC filter, the double monolayer vesicle can be seen in the tunic. (E) Hoechst staining. (E’) Acetylated tubulin counter-staining. (E’’) Composite. (F-I) Screenshots from Supplementary Movie 7 showing tissue left-over fusing with the vasculature at the double vesicle stage, from (F) 0hpi to (G) 5hpi, (H) 10hpi, and (I) 22hpi. (J-K’’) Microscopic view of the areas squared in (I). (J) Transmitted light with DIC filter, the double monolayer vesicle can be seen in the tunic. (K) Hoechst staining. (K’) Acetylated tubulin counter-staining. (K’’) Composite, the insert is a magnification of the region of fusion. (L-O) Screenshots from Supplementary Movie 8 showing tissue left-over fusing with the vasculature at the double vesicle stage, from (L) 0hpi to (M) 7hpi, (N) 20hpi, and (O) 40hpi. (J-K’’) Microscopic view of the areas squared in (O). (P) Transmitted light, the double monolayer vesicle can be seen engulfed by the vasculature. (Q) Hoechst staining. (Q’) Acetylated tubulin counter-staining. (Q’’) Composite. (R-T) Screenshots from Supplementary Movie 9 showing tissue left-over fusing with the vasculature from (R) 0hpi to (S) 19hpi and (T) 22hpi. (U-U’’) Microscopic view of the areas squared in (T). (U) Hoechst staining. (U’) Acetylated tubulin counter-staining. (U’’) Orthogonal projections of the confocal stack show the histological continuity between the bud epithelium and the vascular wall. White arrowheads: tissue left-over, red arrowheads: ampullae fusing with the bud, yellow arrowheads: fusion between bud and vascular epithelium, asterisk: neighboring ampulla.

The localization of potential sites of vascular budding was also monitored *a posteriori*, i.e. by direct detection of clusters of cells in dissected colonies without the tracking via the corresponding movie. By screening different genotypes the first visible signs of putative WBR (n=41 different colonies) were detected between 2 and 5 days after surgery (**Supplementary Table 2**). Also in these screening, we observed different scenarios: the WBR onset was often positioned on the side of the colony facing the surgery (internal side of the system), either in a protrusion of the peripheral vessel (**Supplementary Figure 5A**) or inside an ampullae (in 40 of the 41 colonies) (**Supplementary Figure 5B**). In some cases, up to a dozen of hollow vesicles were observed in a single regenerating colony (**Supplementary Figure 5C**) with several of them being present in the same ampulla or vessel outgrowth (**Supplementary Figure 5D-E**). The presence of more complex epithelial structures was also detected (**Supplementary Figure 5F**).

### 3.3 Reconstruction of the early ontogenesis of the intravascular bud onset

To better describe the morphology of the onset of WBR upon injury, as well as to infer the ontogeny of the process, we further described over a hundred (n=109) proliferating intravascular cell clusters detected within the first 3 days after microsurgery. We coupled previously reported observations (Ricci et al., 2016a) with a higher number of observations and more accurate anatomical descriptions and attempted to assess the dynamics of the vascular bud development. The simplest intravascular structure detected upon microsurgery, and absent in undissected colonies, is a cluster of cells tightly associated with the vascular endothelium. These clusters of between 3 and 8 cells (n=9, size ranging from 12 to 19 µm, average size = 16.1 +/-2.5 µm, **Figure 3A-B**) were found close to the vascular epithelium and proliferated (**Figure 3B**). Immunostaining revealed in the cells of such cluster a consistent localization of gamma-tubulin and PKCξ, suggesting that cells within the cluster have an apicobasal polarity (Parker et al., 2013) (**Figure 3C-D**). The size and the number of cells drew us to consider this intravascular structure a putative initial stage of WBR via vascular budding. The detection, very close to the vascular epithelia, of bigger spherical cell clusters (from 6 to 20 cells, n= 28; size ranging from 12 to 28 µm, average size = 16.8 +/-4.3 µm) without a visible lumen, suggest a possible successive stage (**Figure 3E-F**). In larger vascular buds (n=16; size ranging from 23 to 37 µm, average size = 29.6 +/-4.3 µm), a lumen was detected in the center of the vesicle. These buds consisted of a spherical, hollow, monolayered epithelium. The apical localization of PKCξ, and the presence of cilia in the vascular bud cells, showed an epithelialization and cell polarization in the vascular buds (**Figure 3G, H)**. The cells of the bud are monociliated, with their apical membranes facing the bud lumen and associated with tight junctions (**Figure 3I-J**). On the opposite side, facing the vessel lumen, a basal lamina covered the cells and showed a particular thickening when close to the vascular endothelium (**Figure 3K**). Mesenchymal cells, i.e. hemocytes, inside the vesicle.

**Figure 3.**
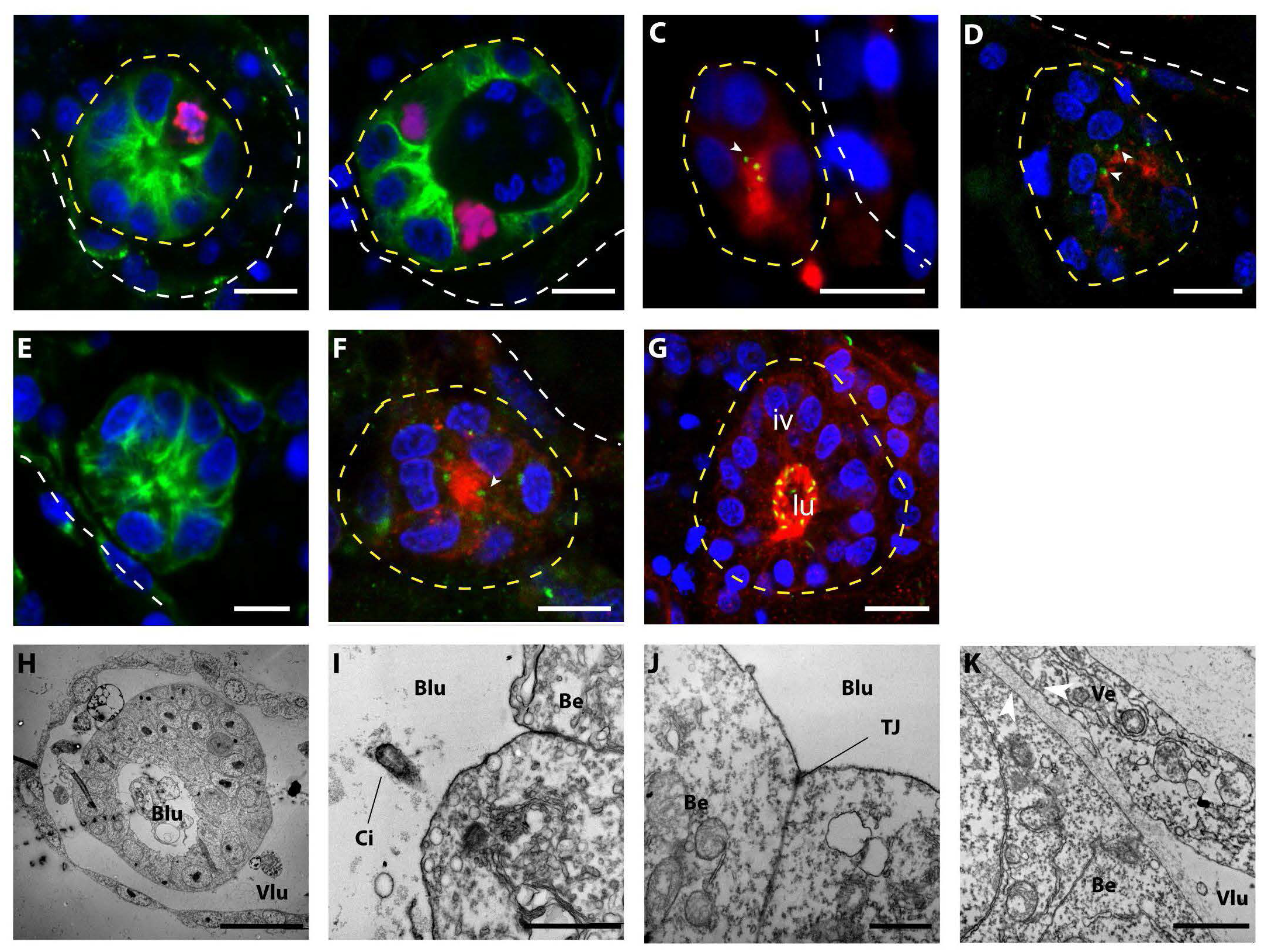
Morphology of small (<40μ) intravascular cell clusters. (A-G) Confocal images of intravascular cell clusters observed 72h post-surgery. The white dotted line points out the epithelia of the vasculature, while the yellow dotted lines highlighted the intravascular cell clusters. Cell nuclei are stained with Hoescht (blue); in (A, B, E) anti-tyrosinated tubulin (green) outlines the cell bodies; in (C, D, F, G) anti-PKCξ (red) shows the apicobasal cell polarity and anti-gamma tubulin (green) suggest the presence of cilia. Iv: inner vesicle, lu: lumen. Scale bar 10μ. (H-K) TEM imaging shows the ultra-structure of the vascular epithelium and the intravascular cell clusters. (H) Overview of an intravascular vesicle lining on the vessel wall; scale bar 20μ.. (H, I) detail of the epithelial cells of the vascular bud, showing monociliated cells (I, scale bar: 1μ) and tight junction (J, scale bar: 1μ) on the apical side (directed towards the bud lumen). (K) Detail of the intracellular cluster and the vessel epithelia. Note the thickening of the basal laminas between vessel epithelial cells and vascular bud cells (arrowheads), scale bar: 1μ. Blu: bud lumen; Vlu: vessel lumen; Ci: cilia; Be: bud epithelium; TJ: tight-junction.

Polarization of the whole vesicle could be observed in the majority of vascular buds of slightly bigger size (n= 37, size ranging from 23 to 55 µm, average size = 41.2 +/- 7.4 µm) (**Figure 4A-C**). In these buds, the side of the bud epithelium in close contact with the vascular endothelium (proximal side) exhibited big, cuboidal cells, with nuclei positioned on the side of the basal membrane. On the opposite side of the bud (distal side), facing the vessel lumen, cells appeared flattened, slightly bigger than their nuclei (**Figure 4A-C**). We also detected a polarized expression of Wnt2 (**Figure 4D**), which is also a marker of polarization in the peribranchial bud (Di Maio et al., 2015).

**Figure 4.**
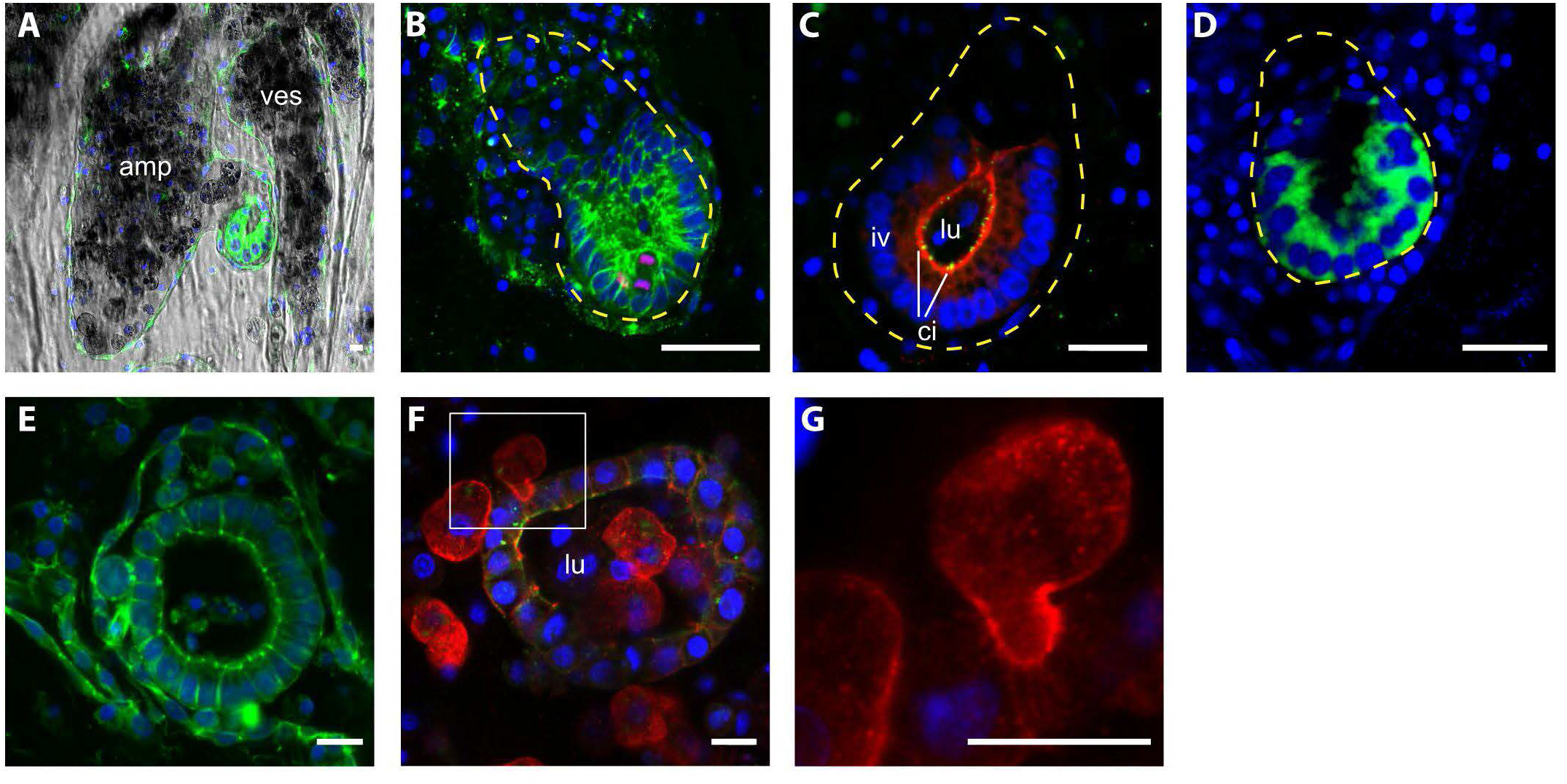
Morphology of intravascular double vesicles. 1 (A) The putative vascular bud grows in the protrusion of an ampulla and (B) it closely interacts with the vascular epithelia, the cells in contact with the vascular epithelia are thicker than the most distal cells. Cell shape is labeled with anti-tyrosinated tubulin (green) and proliferating cells are labeled with anti phospho HH3 (red). (C) Anti-PKCξ (red) shows the apicobasal cell polarity and anti-gamma tubulin (green) shows the presence of cilia. (D) The polarization of the intravascular vesicle is also highlighted by the transient localization of Wnt2 (green). (E-F) Bigger vesicle within the lumen mesenchymal cells, anti-tyrosinated tubulin (green), anti-pan-tyrosine kinase (red). (G) Details of a mesenchymal cell interacting with the epithelia of the vesicle. Cell nuclei are counterstained with Hoescht (blue). Amp: ampullae; ves: vessel; lu: lumen; iv: inner vesicle; ci: cilia. Scale bar: 10μ.

Vesicles with even bigger size, yet without clear polarization were also found (n= 19; size ranging from 45 to 91 µm, average size = 62.6 +/-13.5 µm) (**Figure 4E**). In these latter structures, mesenchymal cells are recurrently present inside the lumen. The position of some mesenchymal cells and their surface activity suggest a dynamic interaction with the vesicle (**Figure 4G-H**).

### 3.4 Inconstant morphogenesis following the double-vesicle stage

After the recurrent scenarios described above, larger vascular buds detected over 3 days post dissections exhibited epithelial folds and compartmentalization of inner cavities, similarly to morphogenesis of peribranchial buds (Manni et al., 2014), although with a greater diversity of configurations of shapes, as suggested by Voskoboynik et al. 2007 (Voskoboynik et al., 2007). While growing, the epithelium of the vascular buds takes the shape of the surrounding vessels and ampullae, resulting in buds distributed in different vascular compartments with completely aberrant forms when compared to blastogenic buds of the same size, including double-axis, *situs inversus,* or hyperplasias (**Supplementary Figure 6**).

### 3.5 Epithelia of the vessels do not contribute to the VB early ontogenesis

To test the possible contribution of the vascular epithelia to the bud onset we took advantage of previous studies that showed the affinity of *B. schlosseri* vascular epithelia for BSA (Braden et al., 2014; Rodriguez et al., 2017, 2018). First, to confirm the specificity of BSA to epithelial versus mesenchymal cells, uninjured colonies were injected with fluorescent-conjugated BSA and counterstained with Hoechst (nuclei) and FRM4-64 (cell walls). Within the first 48h, the presence of BSA is almost exclusively detected in vacuoles inside epithelial cells (98.62% ± 2.38, n= 10, (**Supplementary Figure 7**). After triggering WBR in 12 colonies, 40 VB onsets have been examined at different stages. In none of the vesicles the BSA signal has been detected (**Figure 5**). In 12 cases, BSA was detected in a cell that bridges the epithelial of the vessel with the inner vesicle (**Supplementary Figure 8**).

**Figure 5.**
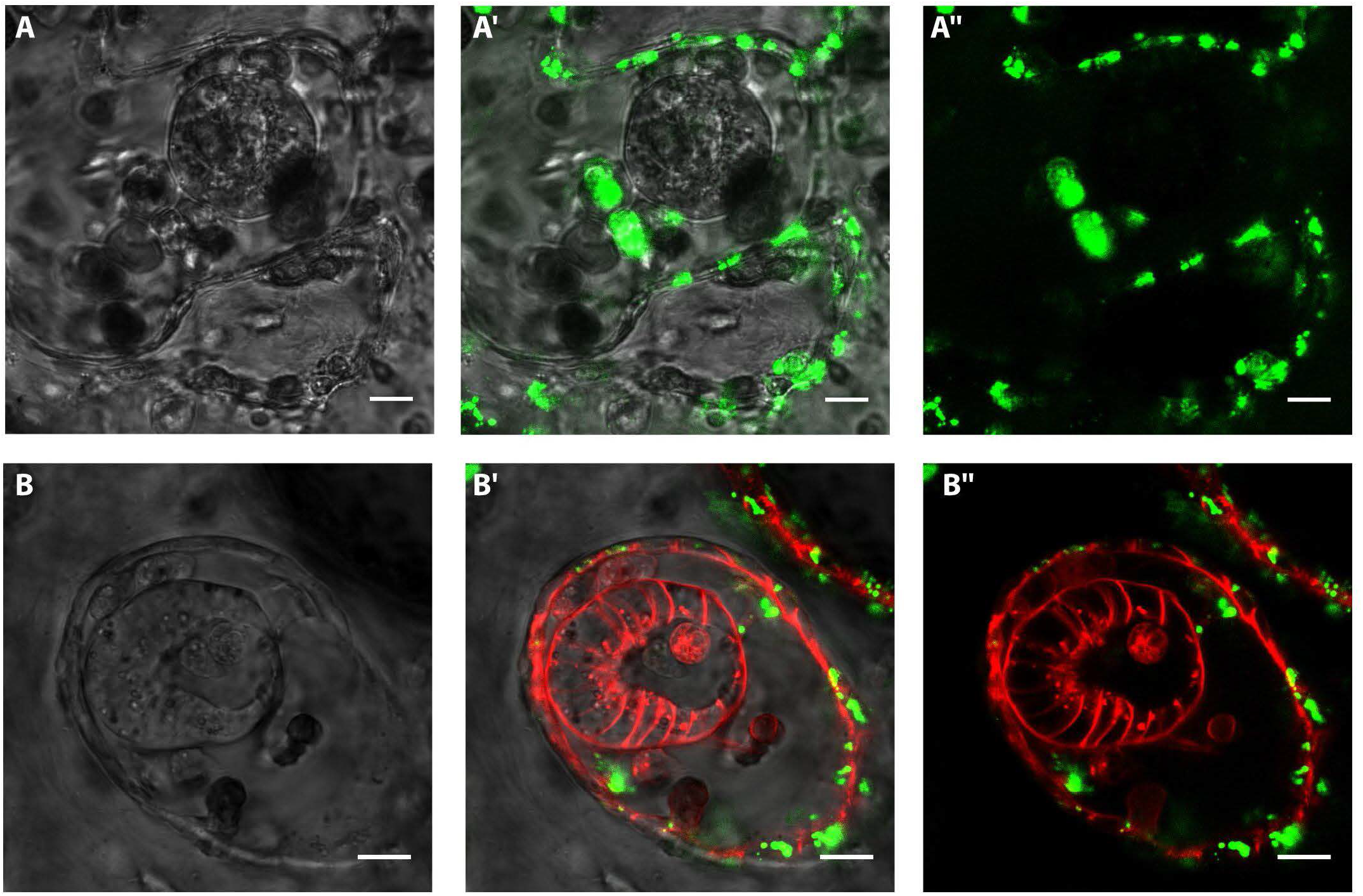
*In vivo* labeling of vasculature epithelium and intravascular vesicles with BSA and FMR4-64. Confocal images of vascular buds after injection of BSA and FM4-64. Green: BSA Alexa Fluor 488 conjugate; red: FM4-64. (A-A’’) Confocal images of a spherical-shaped cluster of cells detected within 3 days after microsurgery. The vasculature is labeled with FITC-conjugated BSA (green). (B-B ’’) Confocal images of a polarized vesicle detected within 3 days after microsurgery. The vasculature is labeled with FITC-conjugated BSA (green), the cell membranes are labeled with FM4-64 (red). Scale bar= 10μ.

### 3.6 In chimeric colonies, regenerating zooids preserve the genotype of the surrounding tissues

To assess whether VB onset is originating from circulating mesenchymal cells, an approach based on allorecognition and chimerism abilities of *Botryllus schlosseri* has been used (McKitrick and De Tomaso, 2010)(See Mat. & Methods). When two individual *B. schlosseri* colonies come into close contact, the ampullae reach out from each individual and come into contact. If the two colonies are histocompatible, the ampullae will fuse and form a single chimeric colony with a common vasculature. Yet, only hemocytes move from one original colony to the other, while the epithelia of vasculature remain separated (Braden et al., 2014; Taketa and De Tomaso, 2015).

After fusion, WBR was induced by depleting zooids and buds from the entire chimeric colony, and the regenerating zooids developed in separate regions of the vasculature. Once they transformed into adult zooids, the gDNA was extracted from their stomachs and their genotype was essayed by analyzing four microsatellite loci and compared with the genotypes of the vascular regions where they appeared (**Supplementary Figure 3C, Supplementary Figure 9**). A total of 36 fusion experiments were performed and 7 of them produced vascular buds with amplified gDNA. For each microsatellite marker, the alleles of the two parental colonies were identified and compared with those of the regenerated zooids (**Supplementary Figure 10**). For the microsatellite BS811, the alleles were different in the parent colonies (size 249 bp for colony A and 229 bp for colony B), and in six out of seven cases, the alleles amplified in the vascular buds corresponded to the genotype of the vasculature from where they originated (**Figure 6**). For the microsatellite PB41 and PB49, the results obtained have the same content, although showing some contamination of the other parent genotype probably due to blood cells. Taken together, these results suggest a vascular origin of WBR or a preferential association between hemocytes and cells of the vascular system of the same genotype.

**Figure 6.**
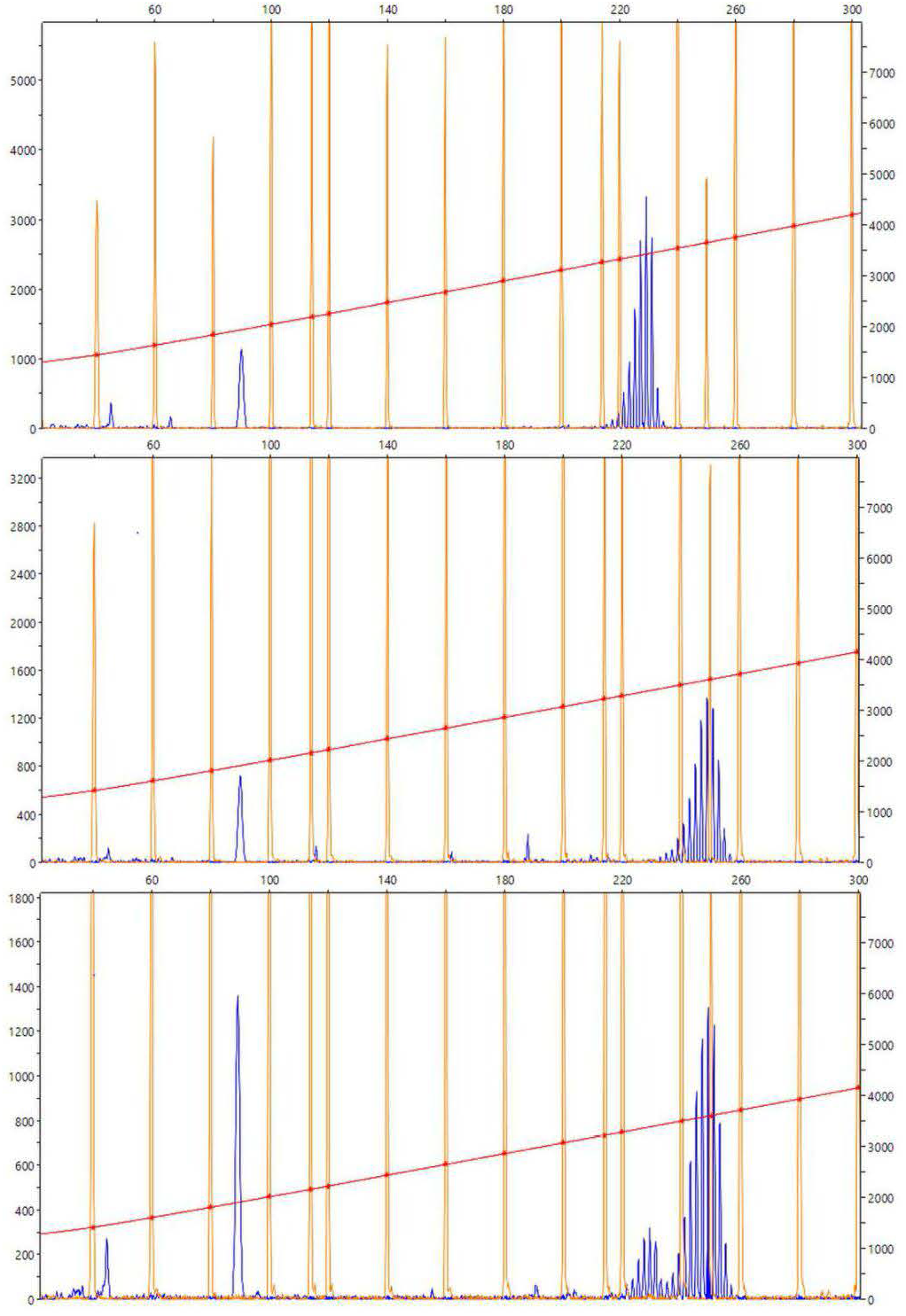
Representative chromatogram for the microsatellites BS811. (A-C) Diagrams showing the size of the BS811 microsatellite locus amplified by PCR. The horizontal axis indicates the size and the vertical axis indicates the intensity of the fluorescence detected in the PCR product. The size is calculated with the default settings of the Peak Scanner software for the referenced standard size used in this experiment, LIZ600 (blue peaks = fluorescence of PCR products; orange peaks = standard size markers). (A) Size of the fluorescence peak detected in the PCR carried out with the gDNA of colonies 1 and 2, collected before the fusion, and with the gDNA obtained from the stomach of a vascular bud, developed in the colony vascular system 2. The size is given in nucleotides. The peak at less than 100 bp could indicate other alleles for the same locus, but since the size of this microsatellite is normally between 200 and 300 bp, it is most likely a non-specific amplification product.

### 3.7 Hemocytes and hemoblasts proliferating activity is stable throughout the VB onset

To provide an overview of the dynamic of cell proliferation after the induction WBR, a time course of their mitotic activity was measured in the early phase of regeneration. While the distribution of mitotic cells appeared scattered through the whole colony all along the time course, since the VB has been detected within ampullae rather than along the vessels, the number of mitotic cells was counted within ampullae of identical volumes at 5 different time points upon injury. By analyzing 89 ampullae of approximately identical volume (1,27∗10^6^±0,06∗10^6^ microns^3^) from 11 different colonies the mitotic activity was detected mainly among circulating hemocytes and it remains stable with a feeble increment at 72h post-injury (**Figure 7** **A-C**).

**Figure 7.**
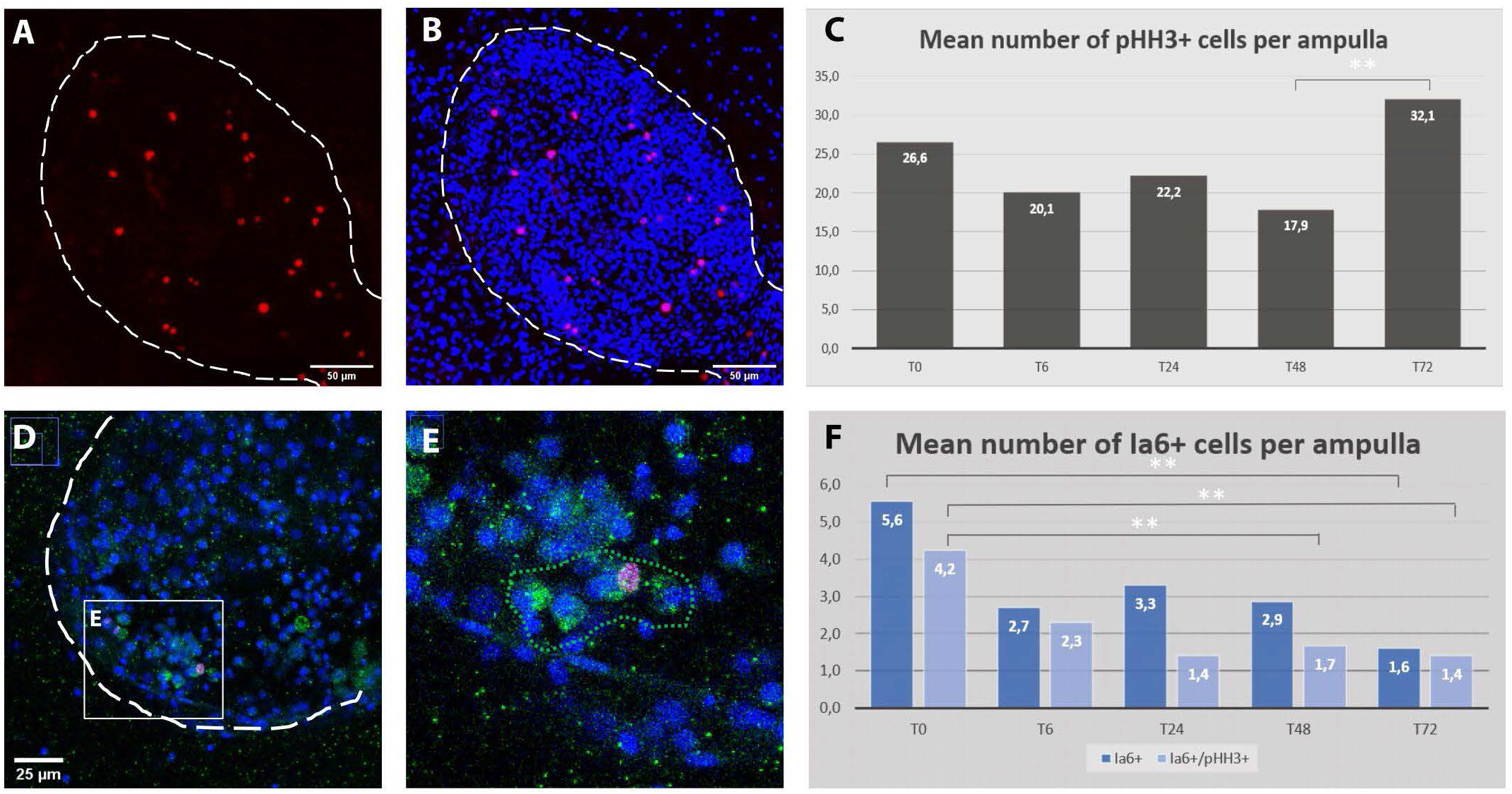
Cell proliferation dynamics and Ia6 expression in ampullae during the first 3 days after microdissection. (A-B) Confocal z-stack showing the detail of an ampulla: proliferating cells are stained with anti-phospho-HH3 (red), cell nuclei are counter-stained with Hoechst (blue), ampulla shape is outlined with dotted lines. (C) Average number of pHH3+ cells per ampulla for five time-points within the first 3 days after microdissection. A significant difference has been detected only between 48 hours (T48) and 72 (T72) hours post microdissection (P=0.025). (D-E) ampulla containing a cluster of Ia6+ cells (green) including one co-labeled with anti-phospho-HH3 (red). (F) Histogram showing the proportion of proliferation Ia6+ cells and their dynamic within the first 3 days after microdissection (P=0.05).

In the closely related species *Botrylloides diegensis,* WBR has been reported to originate from a population of undifferentiated hemocytes, the hemoblasts, which behave like stem cells (Kassmer et al., 2020). Therefore, to explore the behavior of hemoblasts during the early stages of *B.schlosseri* WBR we used the putative hemoblast marker Integrin alpha 6 (Ia6) (Kassmer et al., 2020), assay the presence of Ia6+ and analyzed their dynamics via immunohistochemistry and *in situ* hybridization. With both techniques, we detected the presence of Ia6+. Yet, unlike what has been reported in *B.diegensis,* Ia6+ cells are rare and their number is stable throughout the onset of WBR, decreasing significatively only at 72h post-injury (**Figure 7C-F**, **Supplementary Figure 11**). Similar to *B. diegensis,* the majority of Ia6+ cells are proliferating (**Figure 7C**).

## 4 Discussion

Among the different taxa that acquired WBR, the interest in tunicates regenerative abilities emerged due to their phylogenetic position as the sister group of vertebrates, and also because their regenerative capabilities are plastic within the sub-phylum, i.e. many species regenerate the whole body via asexual budding or upon extensive injury, others have more restrained regenerative potential (Alié et al., 2020; Nydam et al., 2021). Tunicates of the group of *Botryllinae* and in particular *Botryllus schlosseri* have been used for several decades as experimental laboratory models (Manni et al., 2019). The present study discloses previously undescribed dynamics of the phenomenon of injury triggered whole-body regeneration in *B. schlosseri*, and it adds anatomical and molecular elements that serve as a basis for further mechanistic studies in *B. schlosseri* as well as to compare regenerative processes among closely related chordate species.

### 4.1 Transcriptomic response to injury suggests a role of angiogenesis and complement activation in WBR

Regardless of the extent and the nature of the lost part, regenerative response to an injury generally begins with a reparative event, such as wound-healing, followed by the activation of a developmental program that starts with the activation of precursors and eventually the unfolds of new morphogenesis (Carlson, 2007; Tiozzo and Copley, 2015). The overexpression of angiogenesis-related genes and ECM components, together with the extensive vascular remodeling observed during the first 24 hours, point to an active role of the blood vessels in *Botryllus* WBR. In fact similarities in the use of angiogenic factors between vertebrate endothelium and *Botryllus* vascular cells have already been identified (Gasparini et al., 2007, 2014; Tiozzo et al., 2008b; Braden et al., 2014; reviewed in Rodriguez et al., 2019). In the latter, VEGF and VEGFR regulate the active expansion of the vascular network by sprouting angiogenesis, which is key to the expansion of the colony and to maintain a proper connection between zooids (Gasparini et al., 2008, 2014). In addition, the plasticity of the vascular architecture is controlled by the epithelial cells’ ability to synthesize the extracellular tunic (Gasparini et al., 2007) and to regulate its stiffness (Rodriguez et al., 2017). Finally, the ability of ampullae to actively migrate is central in the ability of *Botryllus* to regenerate its vasculature and is controlled by the expression of *BsVEGFR* in epithelial cells (Tiozzo et al., 2008c). Surprisingly, we could not find the *BsVEGFR* transcript in our RNAseq data. However, we found a dynamic expression of several angiogenic factors, of putative growth factors having EGF domains and of many components of the ECM, opening to further functional studies about the role of angiogenesis during WBR in *Botryllus schlosseri*.

Correct regeneration of lost organs in vertebrates necessitates a finely tuned interplay between inflammatory response, neovascularization and ECM remodeling in order to recruit stem/progenitor cells to the regenerative area and to organize the rebuilding tissues (reviewed in Mastellos et al., 2013). For instance, beyond its role as sentinels of immunity, C3 stimulate retina regeneration in chicken and mice (Haynes et al., 2013; Peterson et al., 2021), while in mouse complement proteins regulate wound-healing and angiogenesis in a complex, not fully resolved manner (reviewed in Markiewski et al., 2020). In ascidians, C3 expression has been reported in various epithelial cells and hemocytes of several species (Pinto et al., 2003; Raftos et al., 2004; Giacomelli et al., 2012). In *Ciona intestinalis*, C3 and its putative receptor are expressed by phagocytic amoebocytes (Giacomelli et al., 2012) that show a chemotactic behavior toward sources of synthetic bioactive C3a, suggesting that amoebocytes may be recruited to inflammatory regions (Melillo et al., 2006). In *Botryllus schlosseri*, C3 and components of the lectin pathway are expressed by cytotoxic morula cells that promote phagocytosis of non-self particles (Franchi and Ballarin, 2014; Nicola and Loriano, 2017; Peronato et al., 2020). 2020). Overexpression of C3 orthologue and lectin pathway components (MASP, Ficolin) in the course of WBR in *Botryllus* suggests that the immune role of this pathway is important in the early steps of WBR. It also raises the intriguing possibility that C3 may be used to direct the migration of cells involved in regeneration, for instance, to orientate the vascular ampullae toward the tissue left-over. Finally, the high expression of coagulation-related genes immediately after injury suggests that the complement-coagulation interplay documented in vertebrates may also take place during *Botryllus* WBR to coordinate blood-clotting, defense against pathogen and tissue restoration.

### 4.2 *Origin of WBR in* Botryllus schlosseri

The cellular origin of WBR via vascular budding in *Botryllinae* has been attributed to undifferentiated hemocytes, referred to as hemoblasts, which home to areas of the vasculature and initiate to develop into the regenerating zooid (Rinkevich et al., 1995; Kassmer et al., 2020). The cluster of hemocytes proliferate and differentiate into a hollow monolayered vesicle, which grows in size and gets enclosed by the surrounding vascular epithelia (Brown et al., 2009; Kassmer et al., 2020). The resulting double vesicle is comparable to the one observed during other forms of budding across colonial tunicates, e.g. peribranchial budding in Stolidobranchs (Manni et al., 2014; Ricci et al., 2016a).

In our previous work, Ricci et al. (2016) suggested that in *Botryllus schlosseri* the VB arises from a cluster of mesenchymal cells circulating in the vasculature that gives rise to vesicles eventually developing into a zooid. However, the study lacked longitudinal analyses to backtrack the origin of the clusters, and also missed detailed morphological descriptions to follow the ontogeny of the process (Ricci et al., 2016a). Discordantly, a recently published work by Nourizadeh et al. (2021) proposed that VB originates from, and occurs only, if parts of the blastogenic buds are left behind during the surgery, and therefore suggests an ectopic origin of the WBR. Nevertheless, this study also lacked to follow the *in vivo* dynamics of the process at the cellular level and therefore failed to detect any intravascular vesicle or cell cluster (Nourizadeh et al., 2021).

Indeed, our observations suggest that in *Botryllus schlosseri* vascular buds do not originate from mesenchymal cells resident inside the vasculature but tissues hailed from outside the vasculature and left behind during the injury. In our experimental setup, during the microdissection procedure, the whole blastogenic buds are removed, and so are the majority of the adult zooid tissues. Only small residues of the anterior part of the differentiated adult zooids (<50-70μm) often remain attached to the tunic (**Supplementary Movie 3-5**). According to the anatomy of *B. schlosseri,* these residues may contain parts of the epidermis, the epithelium of the endostyle, the branchial and peribranchial epithelia, and portions of the mantels with residues of muscle fibers and/or peripheral nerves (Tiozzo et al., 2008b; Manni et al., 2014). Starting from this scenario, we observed that within 72h the heterogeneous tissue leftovers: a) migrate and fuse into the vascular network and b) re-shape into different types and numbers of monolayered vesicles (**Figure 8**). First, the migration dynamics potentially involve some form of chemotaxis, which allows the migration of the residual tissues through the tunic and towards the vascular network, as well as angiogenetic/vasculogenetic mechanisms that allow the active sprouting of the tip of the vessels towards the tissue leftover. Second, the reshaping of the tissues into vesicles that eventually gives rise to a complex body suggests the existence of an unforeseen level of tissue plasticity and cell potency. The possible presence in the leftover tissues of residues of endostyle, which has been suggested to be a somatic stem cell niche in *B. schlosseri* (Voskoboynik et al., 2008), may contribute to the initiation of the WBR via vascular budding. On the other hand, mechanisms of cell de-or transdifferentiation, reported in the WBR in other relatively close tunicate species (Kawamura and Fujiwara, 1995; Kawamura et al., 2018), cannot be ruled out. Without a high-resolution method to live-tracking the cells and tissues it was not possible to provide information concerning the exact nature of the left behind tissues and the mechanisms involved. Yet, the lack of a hemocyte proliferation burst following the injury, and the scarce presence of Ia6+ circulating cells, a marker of putative stem cells in the sister species *Botrylloides diegensis* (Kassmer et al., 2020) does not cue toward the presence of mesenchymal stem-cell-based mechanisms in *B. schlosseri*.

**Figure 8.**
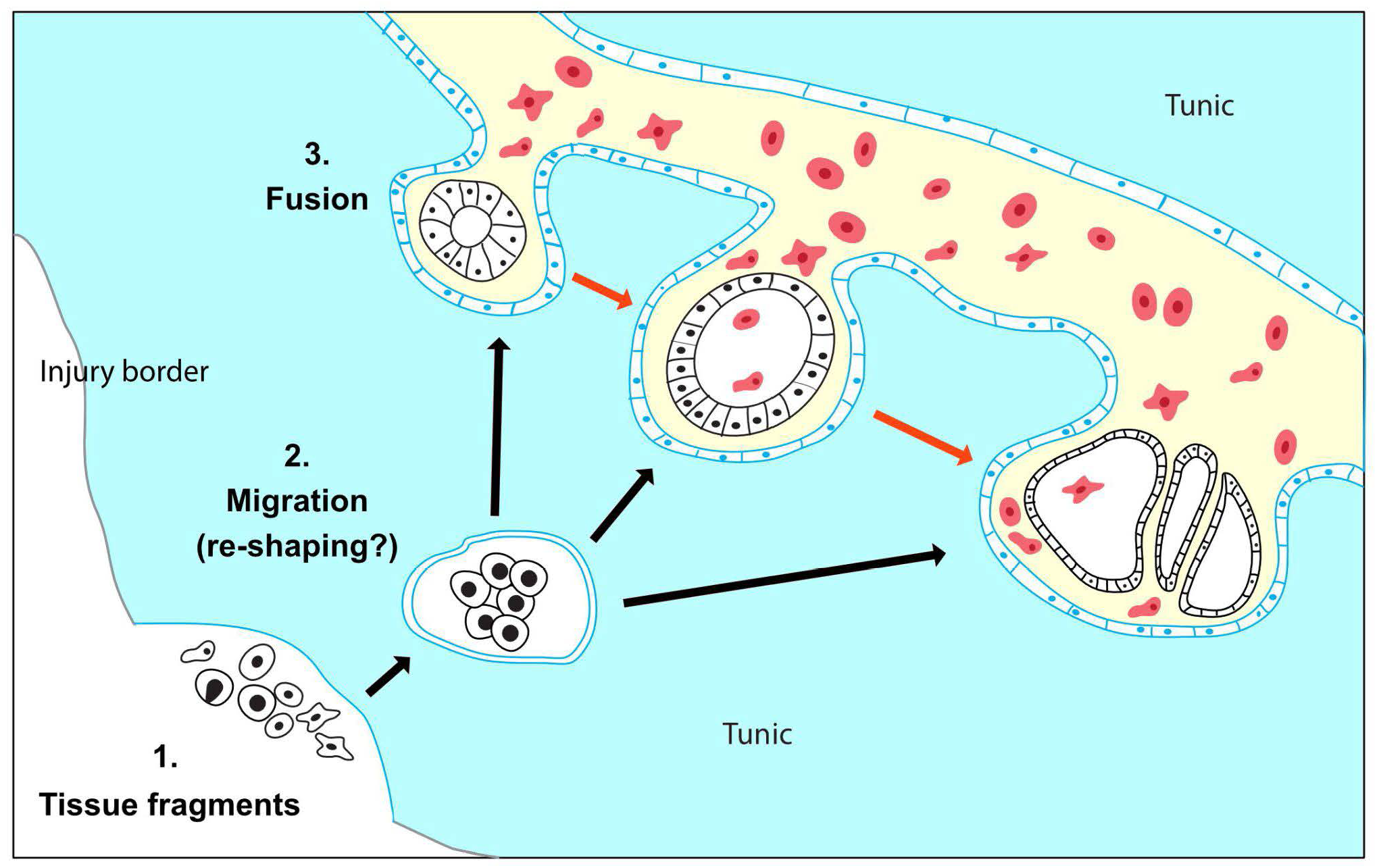
Proposed model for WBR in *Botryllus schlosseri*. (1) WBR originates from heterogeneous tissue fragments of the adult zooid, which have been left behind during injury. (2) The tissue fragments migrate through the tunic (black arrows) and possibly re-shape into spherical vesicles enclosed by a monolayered epithelium (blue). (3) The double vesicles fuse with the vasculature and release the inner vesicle (black) in the vascular system. The vesicles proliferate and develop into the regenerating zooid (red arrows). The exact nature of the leftover cells and the dynamic of tissue reshaping during tissue migration are unknown.

### 4.3 Morphological convergence

The variability of both the site and the time of appearance of the first detectable intravascular structure supports the idea that these two variables are linked respectively to the location and the amount of the tissue left behind upon microdissection. These inconsistencies, which have also been recently reported by Nourizadeh et al. (2021), together with the lack of a proper live-tracking technique do not allow to detail the ontogenesis of the vascular bud once entered into the vasculature. Yet, in our experiments that originally were aimed to completely deplete all zooids and budding tissues, we consistently left behind clusters of 50-70μ circa. In these conditions, the first intravascular structures were observed within a time window of 3 days (72hpi). With the exception of a few cases which showed the presence of complex epithelial structures probably linked to clumsy microdissection (**Figure 2U-U’’**), in most of the microsurgery experiments we detected a variety of intravascular monolayered vesicles. These vesicles were all made of polarized cells, with the apical side facing the lumen. They were all actively proliferating and, when they have been left to develop, they lead to the formation of growing vascular buds. Hence, the lack of contribution of the vascular epithelia and circulating hemoblasts (**Figure 5-6**), the absence of these structures in undissected colonies, and the dynamics seen in the movies strongly suggest that the cellular origin of the vesicle is the tissue leftover derived from the dissected adult.

The monolayered hollow vesicle, which becomes double-vesicle once enveloped by a layer of epithelial tissue, is a phylotypic stage common to many types of budding in tunicates (Alié et al., 2020). We previously documented that at this stage, the regionalized expression of germ-layers markers suggests a cell commitment (Ricci et al., 2016a). Therefore, the regular detection of this structure, its continuous proliferative activity, and the commitment of its cells suggest that a morphogenetic program is already in place. Unlike the vascular budding in other *Botryllinae*, the morphogenesis has been documented to unfold abnormally, regaining the normal developmental patterns only after a series of generations of blastogenic budding (Voskoboynik et al., 2007). While further observations are needed, the abnormalities detected seem to concern the patterning (axes and a/symmetries) rather than the cell differentiation, as the presence of differentiated muscles and the nervous system seems to suggest (**Supplementary Figure 6**).

### 4.4 Variation of injury-induced WBR capacities across *Botryllinae*

Among tunicates, both the diversity of the cellular onsets and the phylogenetic distribution suggests that the WBR capacity via propagative and survival budding is a plastic trait that evolved multiple times (Alié et al., 2018; Nydam, 2020). Mesenchymal stem cell-driven budding like vascular budding, has been suggested as a propagative and/or survival mode of WBR in several species of tunicates and it has been documented in *Botryllus schlosseri’*s closest related species such as *Botrylloides diegensis* (**Supplementary Figure 1**). In the latter, as well in other *Botryllinae* species, VB can be easily induced by isolating a small portion of the extracorporeal vasculature. On the other hand, in *B. schlosseri* a more structured vascular network and the presence of extravascular tissue seems to be necessary for the WBR to start (Sabbadin et al., 1975; Nourizadeh et al., 2021). Therefore, even if we cannot rule out a “leftover-free” initiation of VB, all the data collected seems to suggest that *Botryllus schlosseri* does not undergo vascular budding as the other *Botryllinae*. Such phylogenetic proximity offers the opportunity to identify at the intra-generic level the genomic basis of developmental plasticity linked to whole-body regeneration.

## 5 Data availability statement

The transcriptomic datasets generated in this study will be available upon request and deposited in the Gene Expression Omnibus repository after acceptance of the manuscript.

## 6 Author contributions

ST and LR designed the study; LR, BS, CO, AA, and ST performed the experiments; AA, RAP, and AC assembled the transcriptomes and analyzed the RNAseq dataset, ST and AA wrote and edited the manuscript. All authors read and commented on drafts, and approved the final version.

## Acknowledgments

We would like to thank the EMBRC-France and in particular Laurent Gilletta for setting up the aquaculture system, Mohamed Khamla for help with the video editing, Sonia Lotito for the technical support, Laurel Hiebert and Philippe Dru for help with the bioinformatics resources.

## Figures

**Supplementary Figure 1.**
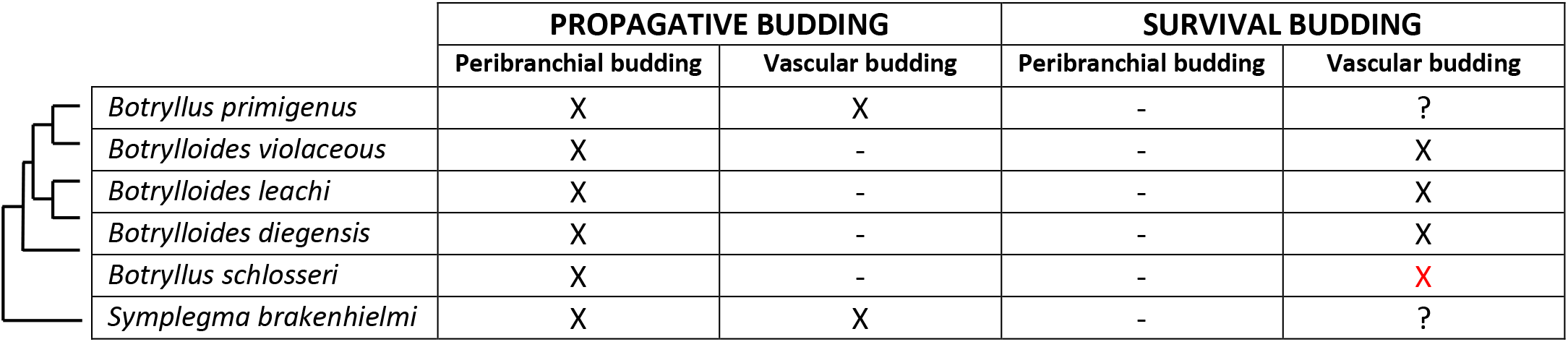
Mode of budding in the five species of Botryllidae. The budding has been classified according to Nakauchi (1982) as part of the asexual life-cycle (propagative) or as a response to injury (survival), as well as according to the tissue origin and ontogenesis. The phylogenetic relationships among species have been obtained from Salonna et al. 2021. The presence of survival vascular budding in *B.schlosseri* (red X) is discussed in the present manuscript.

**Supplementary figure 2.**
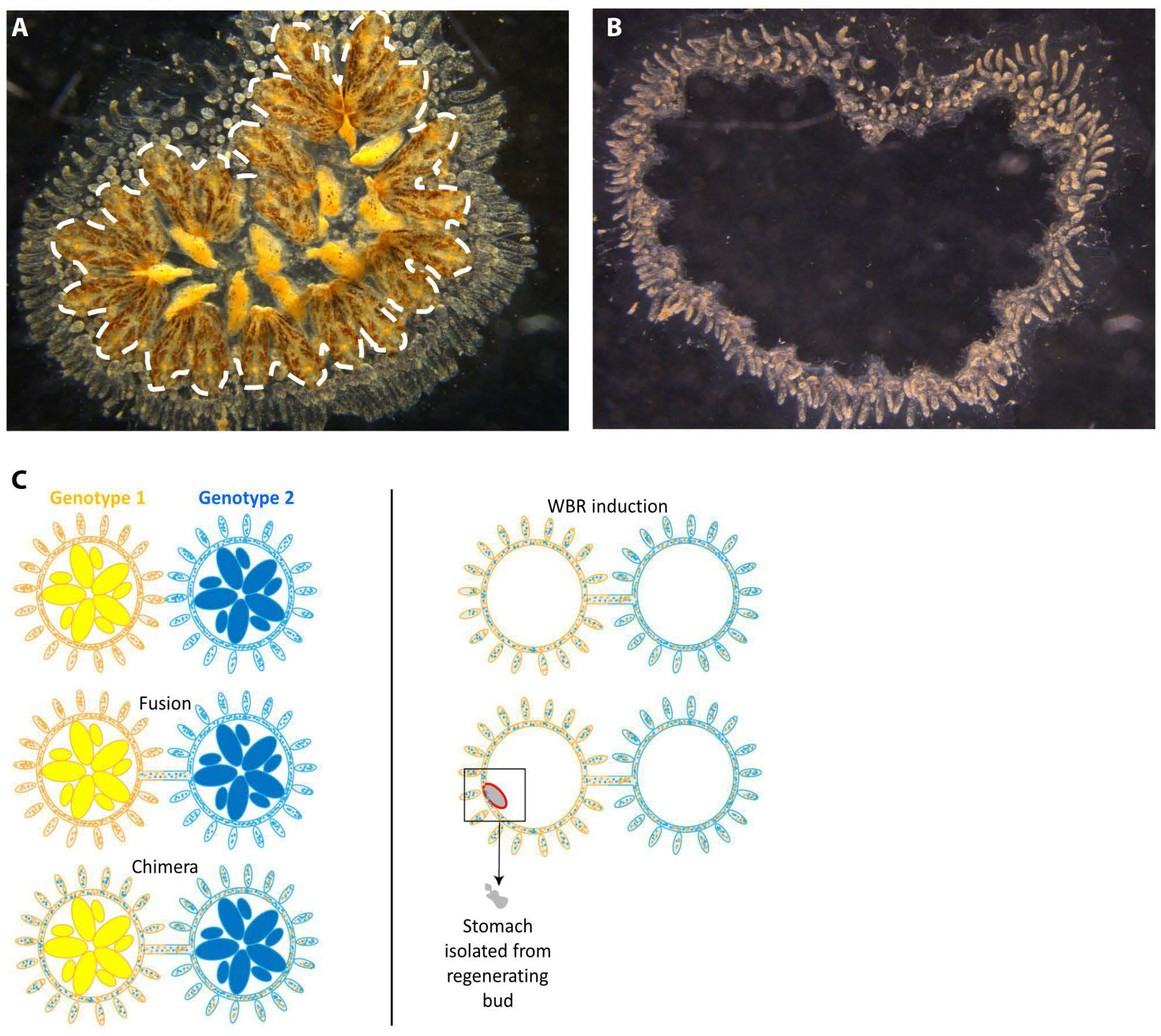
Induction of WBR in *Botryllus schlosseri* and experimental design of the induction of WBR in chimeras for the genotyping of vascular buds. (A-B) Dorsal view of a *B. schlosseri* colony (A) before and (B) after microsurgery. (C) Diagrams experiment of chimerism and microsurgery. Colors rapresent the two different genotypes. The colored dots rapresents the hemocytes. Colonies corresponding to genotype 1 and 2 have been previously assaied for histocompatibility.

WNT2

**See:** (Di Maio et al., 2015)

Di Maio, A., Setar, L., Tiozzo, S., and De Tomaso, A. W. (2015). Wnt affects symmetry and morphogenesis during post-embryonic development in colonial chordates. *Evodevo* 6, 17. doi: 10.1186/s13227-015-0009-3.

INTEGRIN ALPHA-6

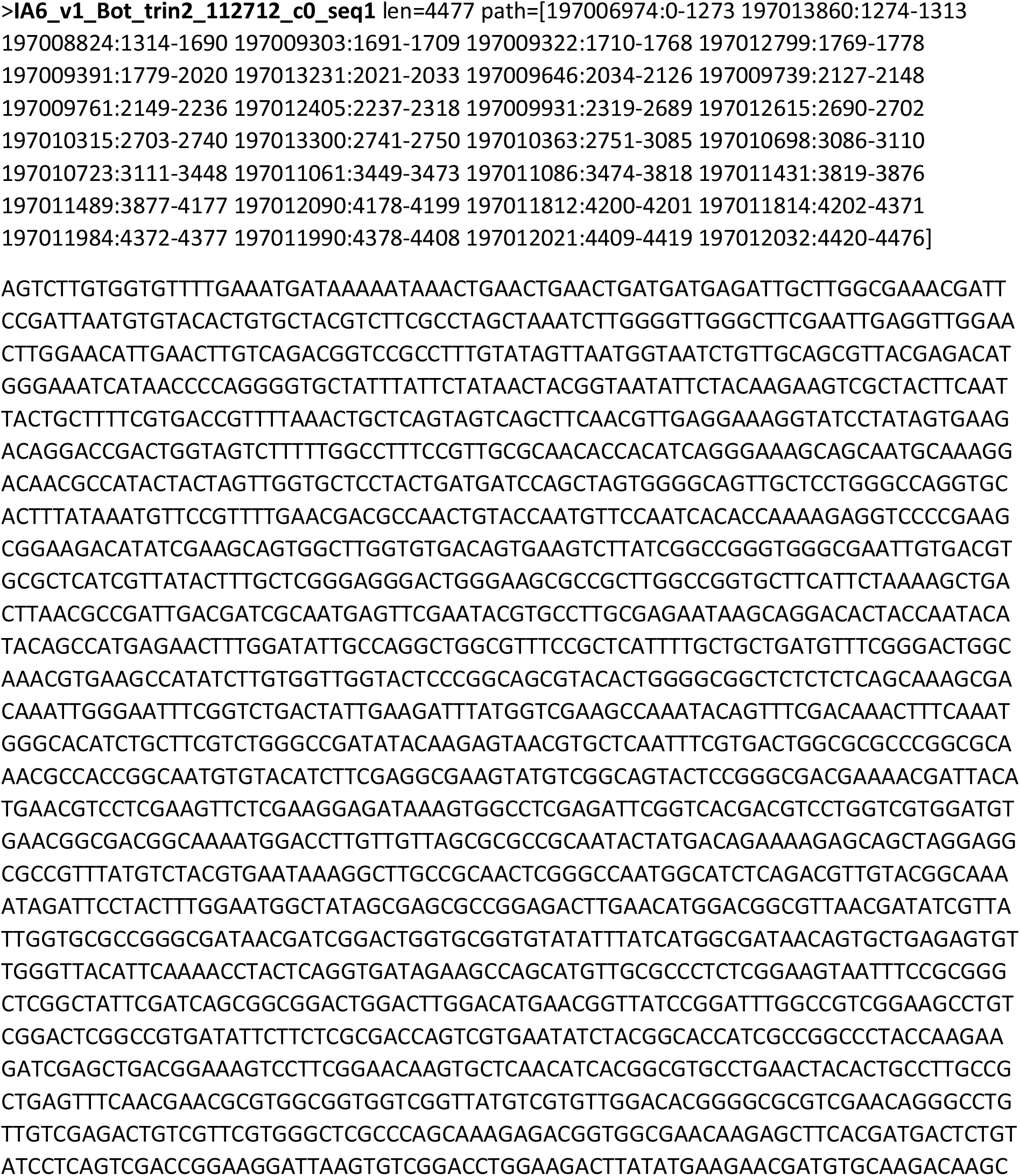
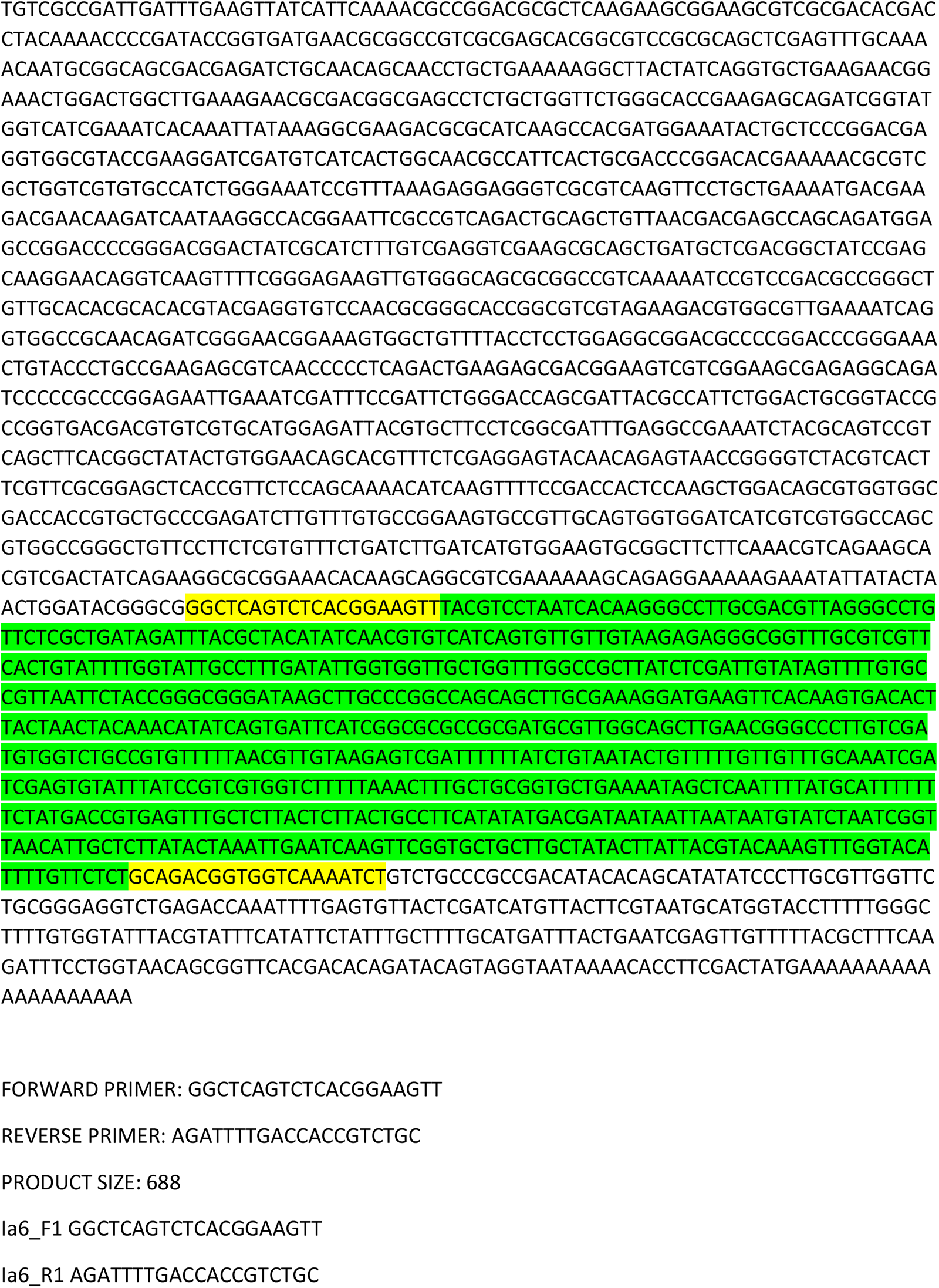

VASA

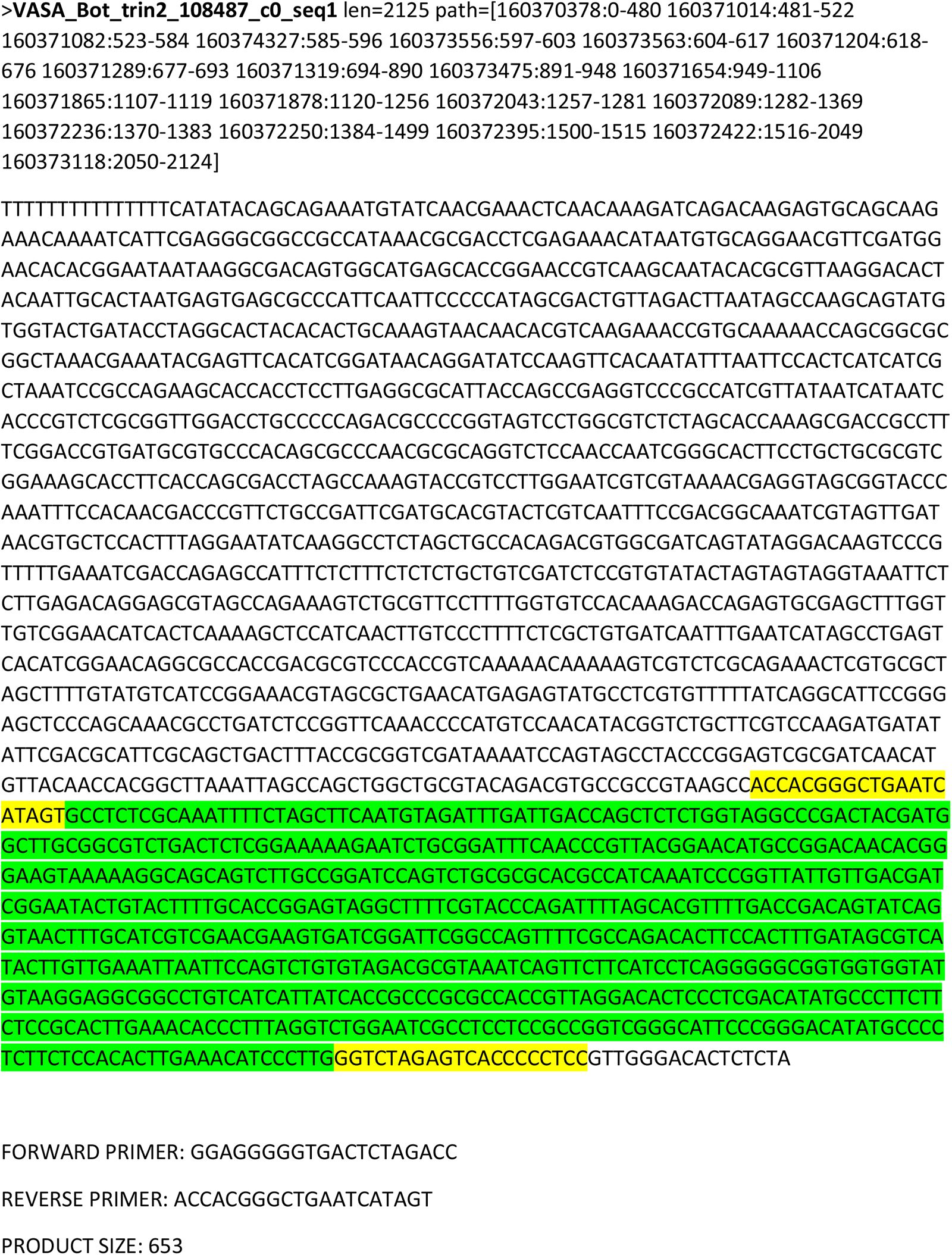

**Supplementary Figure 4.**
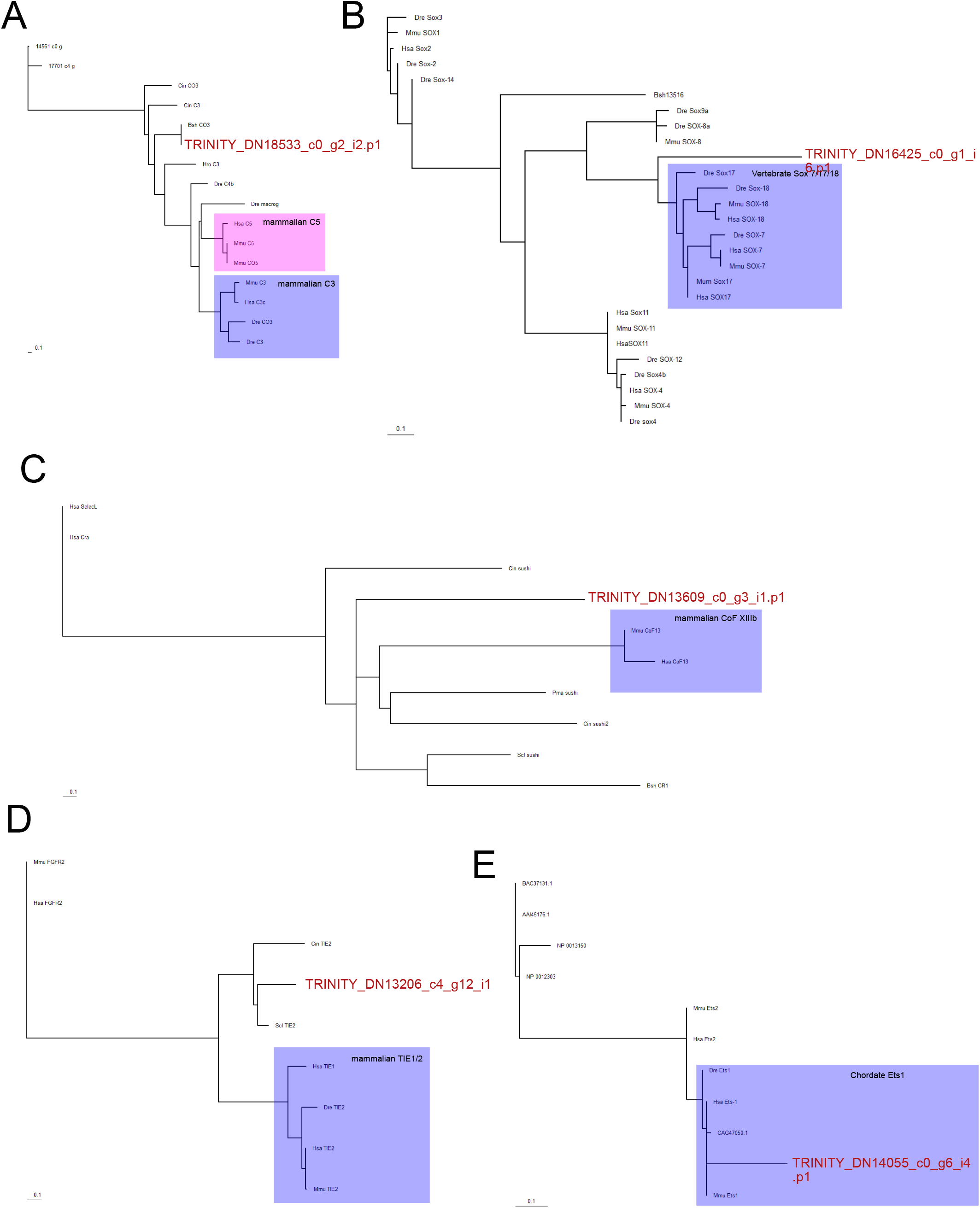
Phylogenetic analyses of genes cited in the results. (A) Complement C3/5. (B) Sox 7/17/18. (C) Coagulation Factor XIIIb. (D) TIE1/2. (E) Ets-1. Maximum likelihood analyses using PhyML, model LG +G, NNI/SPR mix search. Tree were visually rooted on vertebrate putative paraphyletic outgroup based on blast results. Abbreviations : Bsh, *Botryllus schlosseri*; Cin, *Ciona intestinalis*; Dre, *Danio rerio*; Hro, *Halocynthia roretzi*; Mmu, *Mus musculus*; Hsa, *Homo sapiens*; Pma, *Phallusia mammillata*; Scl, *Styela clava.* In red : contigs of *Botryllus schlosseri*.

**Supplementary figure 5.**
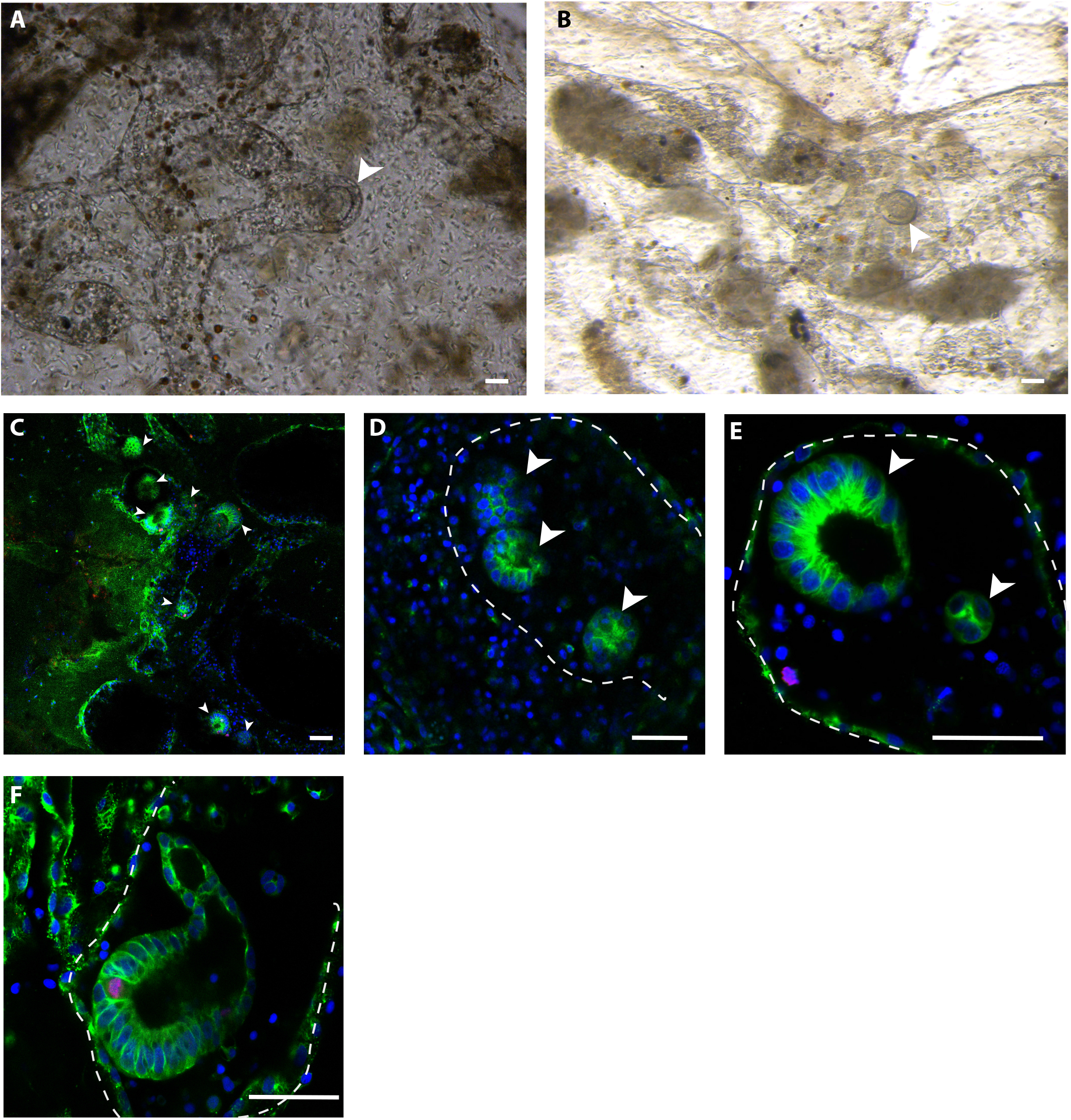
(A-B) *In vivo* early detection of intravascular vesicles observed withing 3 days after microsurgery. (C-D) Confocal images showing the different structures detected inside the vasculature withing 3 days upon microsurgery, cell shapes are labelled with anti-tyrosinated tubulin (green), proliferating cells are labelled with anti phospho HH3 (red) and cell nuclei are counterstained with Hoescht (blue). (C) presence of numerous monolayered vesicles (arrowheads); (D) details of one ampullae with three monolayered vesicles (arrowheads); (E) detail of polarized monolayered vesicele and cluster of cells (arrowheads) in the tip of a vessel; (F) presence of more complex epithelial structures inside an ampullae. White dotted-line highlight the epithelia of the vessel. Scale bar: 10μ

**Supplementary figure 6.**
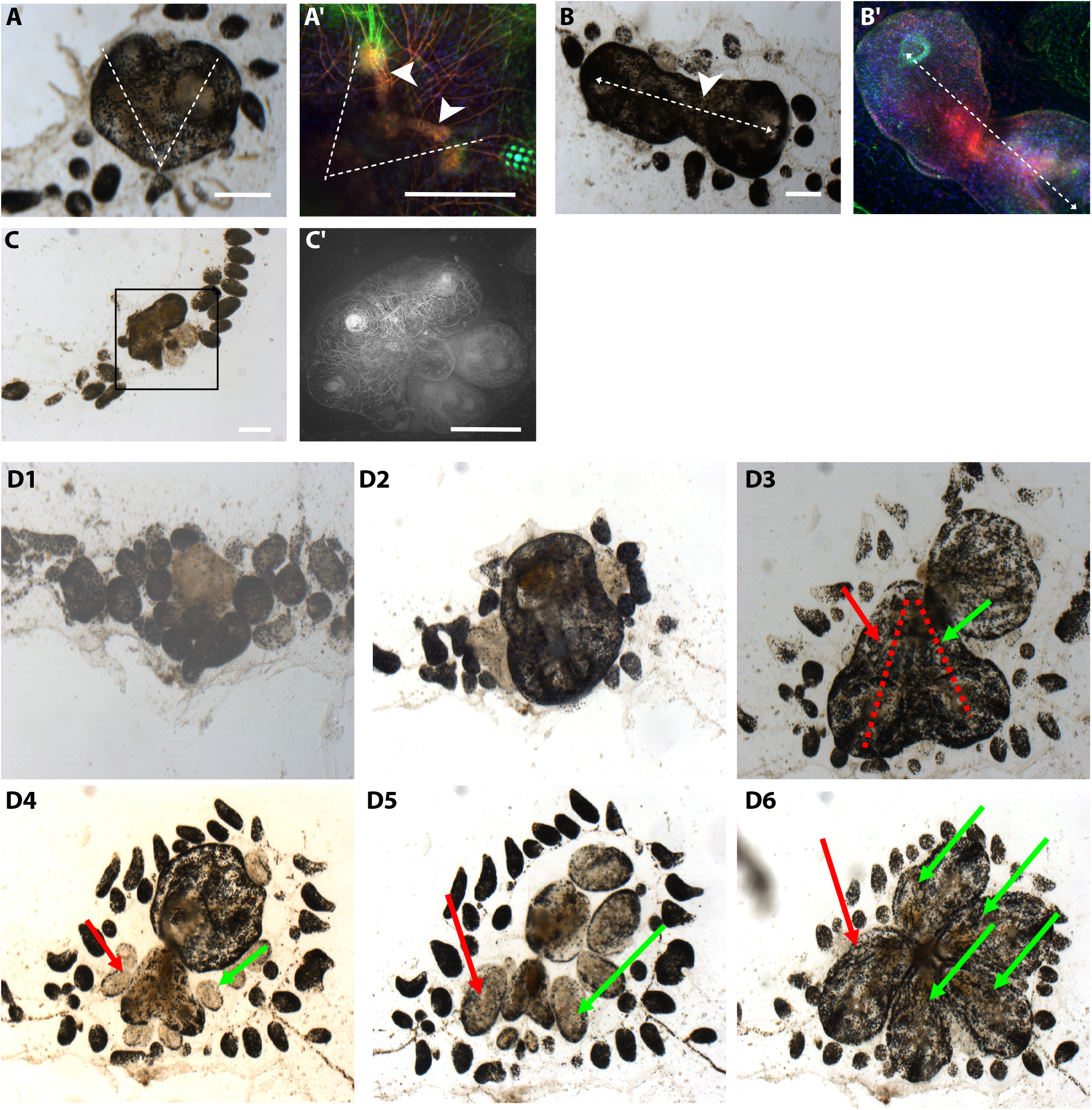
(A) *In vivo* image of abnormal zooid showing duplication of antero-posterior axis (dotted lines). (A’) Confocal image showing the detail of the duplication of the neural gland and ciliated funnel (arrowheads),anti-acetylated tubulin (green), phalloidin (red) and cell nuclei are counterstained with Hoescht (blue). (B) *In vivo* image of abnormal zooid showing posterior fusion (arrowhead) along the anto-posterior axis. (B’) Confocal image showing the same abnormal zooid labelled with anti-acetylated tubulin (green), phalloidin (red) and Hoescht (blue). (C) *In vivo* image of abnormal fused zooids. (C’) Confocal image showing the same abnormal zooid labelled with phalloidin. (D1-D6) Growth of a vascular bud, followed by asexual budding. (D3) Abnormal zooid showing anterior-posterior axis duplication (red dotted lines): the duplicated hearts (arrows) are inverted. The red arrow shows the correct location of the heart (right side of the zooid) and the green arrows shows the hearts located in the left side of the zooid. Scale bar: 100μ.

**Supplementary figure 7.**
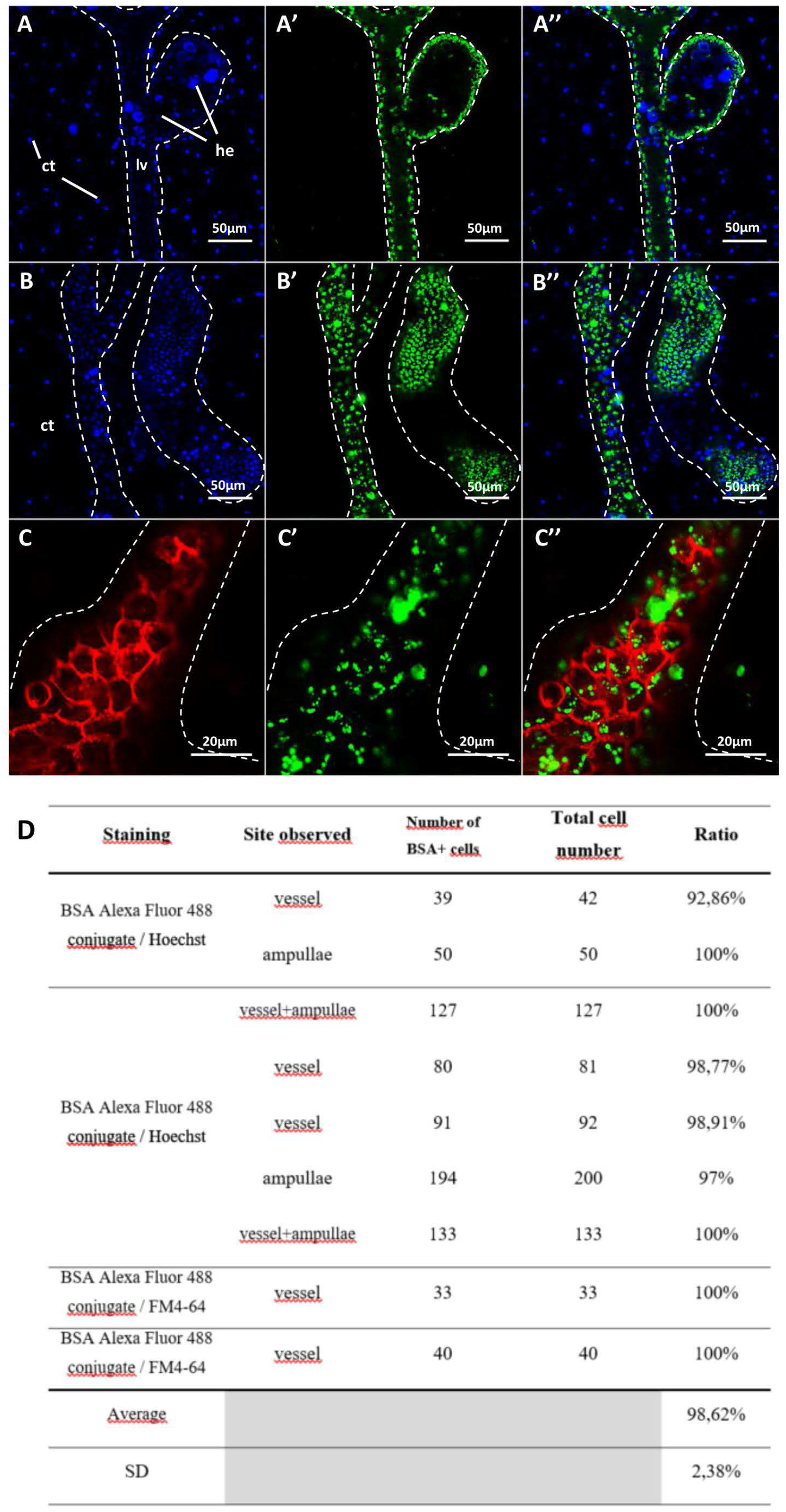
Specific *in vivo* labeling of vascular epithelium with fluorescent BSA. Confocal images of vessels and ampullae labeled with BSA conjugated with Alexa Fluor 488 (green), cell nuclei are counter stained with Hoechst (blue). (C-C”) Cell membrane is stained with the vital dye FM4-64 (red). A-A “:vessel and small bulb where 127 nuclei belonging to the vascular walls were counted; B-B “:ampullae and vessel where 133 nuclei were counted; C-C”:top ofvessel where 33 cells were counted, all containing at least one green fluorescent spot. (D) Colonies analized to calculate the ratio ofBSA+ epithelial cells.

**Supplementary figure 8.**
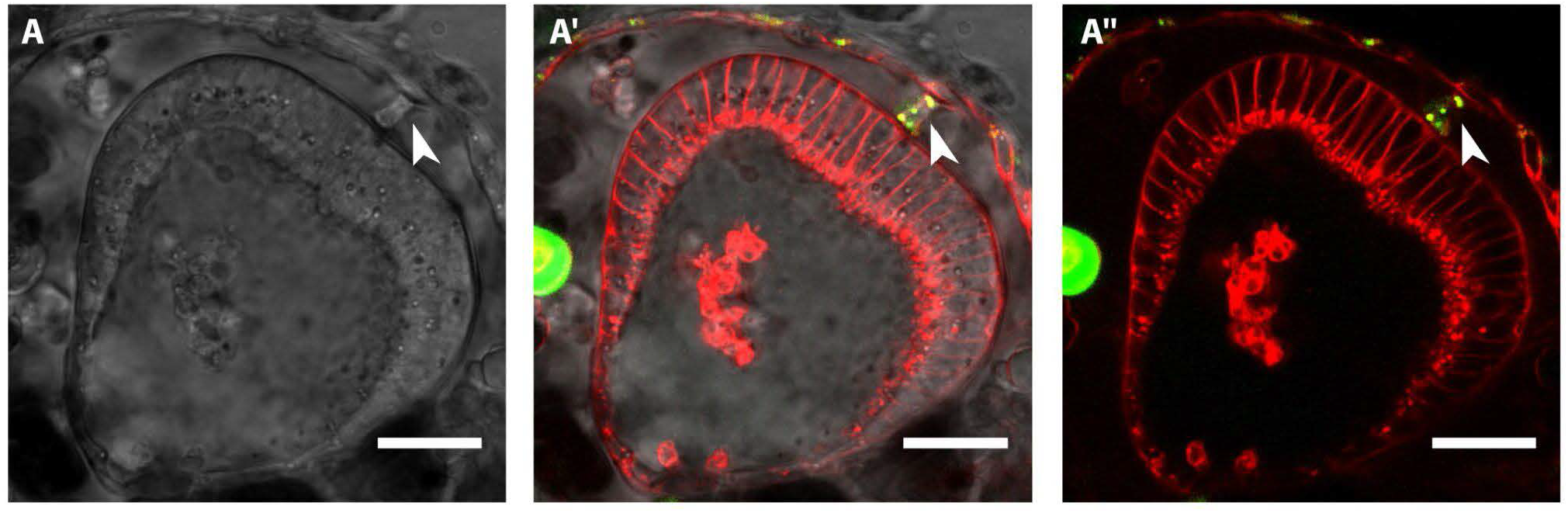
Confocal image showing *in vivo* a polarized intravascular monolayered vesicle. Arrowhead shows a BSA+ cell connecting the epithelial vasculature with the vesicle. BSA conjugated with Alexa Fluor 488 (green), cell membrane is stained with the vital dye FM4-64 (red). Scale bar: 10μ).

**Supplementary figure 9.**
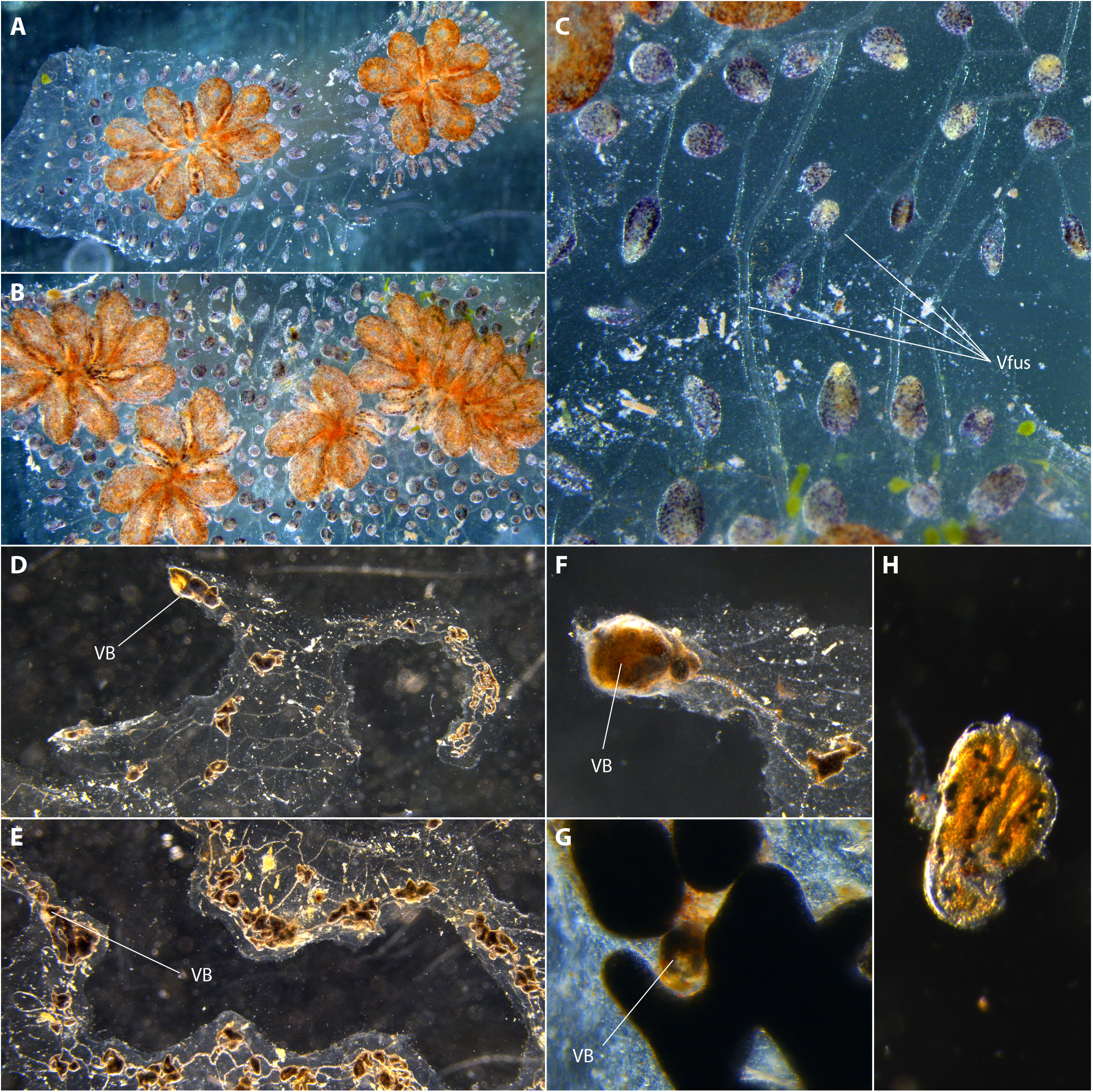
Sampling of vascular buds from regenerating chimeric colony. (A-B): colonies fused before dissection; (C) detail of the site of fusion between the two colonies; (D and E) regenerating zooid at 13 days after microdisection; (F-G): vascular buds 27 days after microdissection; (H) stomach dissected from a vascular bud (27 days after microdissection). VB = vascular bud; Vfus = fused vessels.

**Supplementary Figure 10.**
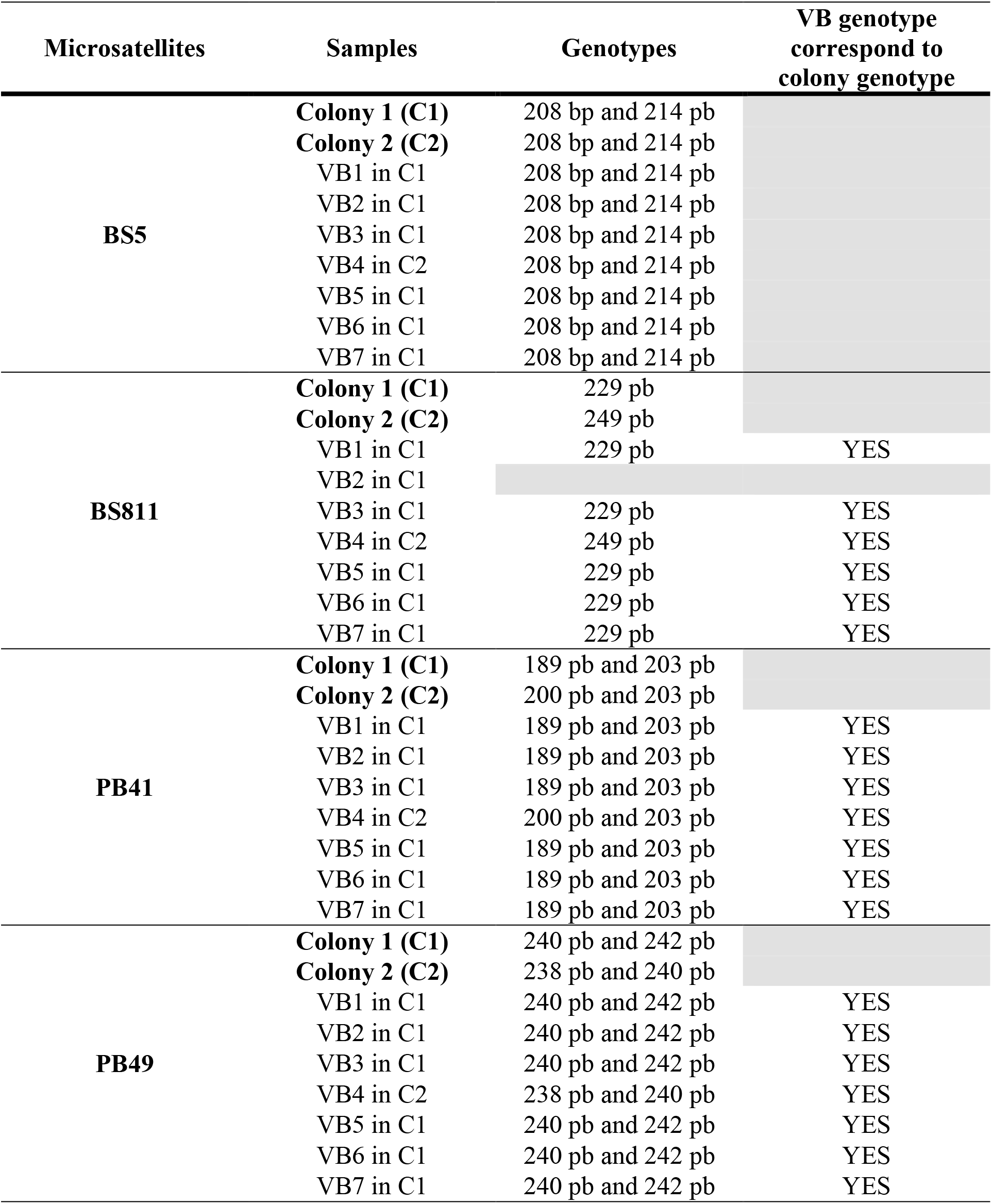
Summary table of microsatellite genotyping results

**Supplementary Figure 11.**
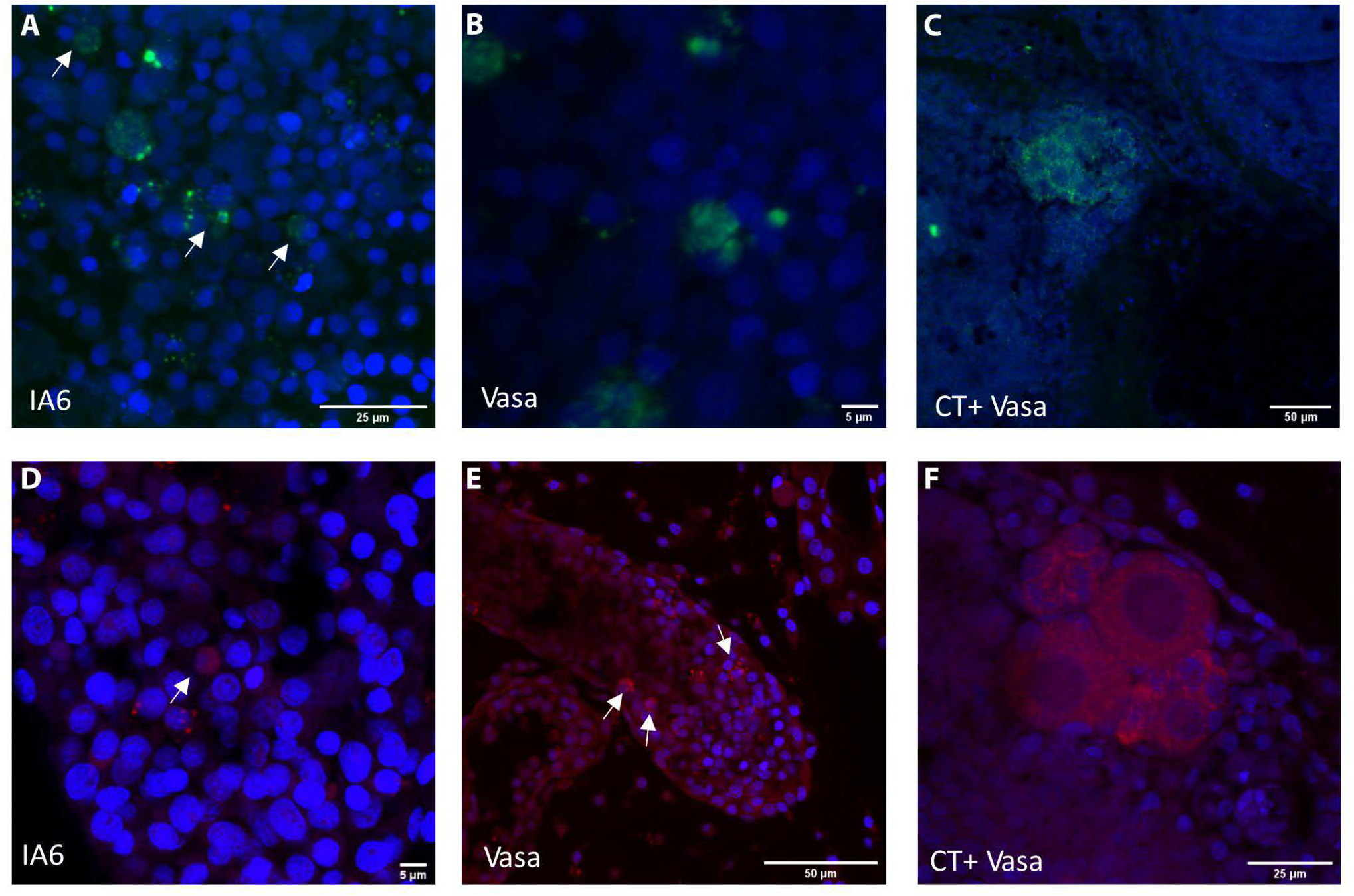
Fluorescent whole-mount in situ hybridizations labelled with green (A-C) and red (D-F) ftuorochromes. Nuclei are counter-stained with Hoescht (blue). (A-D) Confocal stacks of the details of an ampullae showing the presence of Integrin-alpha-6 cells (arrows). (B-E) Confocal stacks of the details of an ampullae showing the presence of Vasa positive cells (arrows, positive control). (C-F) Female gonads labelled with antisense probe for Vasa (positive control).

