## Supplementary table 2 for "The onset of whole-body regeneration in *Botryllus schlosseri*: morphological and molecular characterization"

| Time of vb detection (days post surgery, dps) | Number of colonies | Percentage |
| --- | --- | --- |
| < 2 dps | 5 | 12% |
| 2 – 3 dps | 14 | 34% |
| 3 – 4 dps | 9 | 22% |
| 4 – 5 dps | 7 | 17% |
| > 5 dps | 6 | 15% |

|  | Clone AM | Clone BG | Clone S |
| --- | --- | --- | --- |
|  | 84 | 36 | 72 |
|  | 72 | 60 | 72 |
|  | 48 | 72 | 72 |
|  | 72 | 72 | 60 |
|  |  | 84 | 84 |
|  |  | 84 | 98 |
|  |  |  | 84 |
| Average (hps) | **69,0** | **68,0** | **77,4** |
| S.D (hours) | **15,1** | **18,1** | **12,3** |

Supplementary table 2: Time of detection of the first vascular bud

(A) Data are provided for *B. schlosseri* colonies collected in Villefranche sur mer, n=41. Average time of detection was 77 hps and was calculated using the median value for each class of time as reference (e.g., 14 and 9 vascular buds were detected at 2.5 dps and 3.5 dps, respectively). (B) The table shows the time (in hours post-surgery, hps) of vascular bud first detection for three different genotypes of *Botryllus* colonies: AM, BG and S clones, n= 4, 6 and 7, respectively.
